# Cytosolic MAPK signaling gates chloroplast protein import and photosynthetic capacity

**DOI:** 10.1101/2025.11.22.689956

**Authors:** Sarvesh Jonwal, Balakrishnan Rengasamy, Gopal Banerjee, Muskan Bansal, Mohit Mohit, Pallavi Sharma, Alok Krishna Sinha

## Abstract

Chloroplast protein import is essential for photosynthesis, yet whether and how cytosolic signaling pathways dynamically regulate this process remains largely unknown. Here we uncover a signaling module that links mitogen activated protein kinase (MAPK) activity to fine tune the import of small subunit of Rubisco (RbcS) into the chloroplast. We show that MPK3 negatively regulates the cytosolic Raf-like kinase ACTPK1, which directly phosphorylates the transit peptide of RbcS precursor at Thr12. MPK3 directly phosphorylates ACTPK1, attenuating its kinase activity and thereby limiting RbcS transit peptide phosphorylation. Genetic and physiological analyses demonstrate that loss of MPK3 elevates ACTPK1 activity, increases RbcS phosphorylation and enhances Rubisco accumulation and CO₂ assimilation. On the other hand, ACTPK1 deficiency compromises these processes and photosynthetic performance. Phospho-mutant analyses further reveal that reversible phosphorylation of the RbcS transit peptide is required for efficient chloroplast import. Together, our findings establish chloroplast protein import as a signaling-regulated process and identify transit peptide phosphorylation a key check point integrating cytosolic MAPK signaling with photosynthetic capacity.

## Introduction

Photosynthesis is the key photochemical process that drives the conversion of CO₂ and H₂O into organic carbon compounds. This process is regulated by multiple leaf traits, encompassing physiology, biochemistry, and development along with external environmental factors. Photosynthetic proteins critical to this process are encoded by both the nuclear and chloroplast genomes. Chloroplast-encoded proteins are synthesized within the chloroplast, while nucleus-encoded chloroplastic proteins are transcribed in the nucleus, translated in the cytosol, and translocated to the chloroplast via a targeting signal peptide that directs them through the chloroplast membranes (Bruce, 2000; Jarvis & López-Juez, 2013). Once inside the chloroplast, these proteins are processed and assembled into functional complexes. Rubisco, a pivotal enzyme in CO₂ fixation, is a hexadecameric holoenzyme comprising eight large subunits (RbcL) and eight small subunits (RbcS). The translocation of the small subunit into the chloroplast is essential for holoenzyme assembly. In the cytosol, RbcS is synthesized as a precursor bearing an N-terminal transit peptide, stabilized by chaperones such as Hsp70 and targeted to the chloroplast, where it is processed, folded and assembled with RbcL by stromal chaperones and assembly factors including Cpn60/Cpn10 and RAF1 (Lee et al., 2009, 2013; Jarvis & López-Juez, 2013; Spreitzer, 2003; Feiz et al., 2012; Hauser et al., 2015). Efficient RbcS import is therefore required to stabilize the RbcL octamer and form functional Rubisco, and its failure severely compromises CO₂ fixation and photosynthetic performance (Andersson & Backlund, 2008).

Enhancing leaf photosynthesis is a major goal for improving crop productivity, and biochemical studies have identified multiple strategies to overcome intrinsic limitations of Rubisco and carbon assimilation (Long et al., 2006; Zhu et al., 2010; Gibson et al., 2011; Murchie & Niyogi, 2011; Covshoff & Hibberd, 2012; Ort et al., 2015; Simkin et al., 2019). Photosynthetic capacity is constrained by the catalytic properties of Rubisco, the efficiency of electron transport, RuBP regeneration, and developmental traits that influence light capture and CO₂ diffusion (von Caemmerer, 2000; Parry et al., 2007; Sharkey et al., 2007; Murchie & Lawson, 2013; Terashima et al., 2011; Mathan et al., 2016). While these biochemical and physiological determinants have been extensively characterized, the information about how regulatory signaling pathways control the targeting, stability and accumulation of photosynthetic proteins to coordinate photosynthetic performance is poorly understood (Evans, 2013).

Photosynthetic efficiency is further modulated by environmental fluctuations, such as light, temperature, and CO₂ concentration, which impact Rubisco activity, electron flow, and carbon assimilation (Kalaji et al., 2016; Sharma et al., 2020; Borba et al., 2023; Gao et al., 2024). To maintain optimal photosynthesis under these conditions, plants employ post-translational modifications along with gene expression at transcriptional and translational levels, with phosphorylation playing a central role (Puthiyaveetil et al., 2009, 2010; Schönberg & Baginsky, 2012; Jonwal et al., 2021; Xiong D, 2024). Phosphorylation, mediated by kinases such as STN7, STN8, and cpCK2α, targets core photosystem proteins, light harvesting complexes, plastid transcription factors and Calvin cycle enzymes. They function by modulating their activity, stability, and interactions to balance electron flow, protect against photoinhibition, and optimize CO₂ fixation (Bellafiore et al., 2005; Reiland et al., 2009, 2011; Schönberg et al., 2017). While phosphorylation is a well-established mechanism in photosynthetic regulation, the role of Mitogen-Activated Protein Kinase (MAPK) cascades in this context remains underexplored.

MAPK cascades are conserved signalling pathways in eukaryotes. They operate in three tiers: MAPKKKs activate MAPKKs at a conserved S/T-X5-S/T motif, which in turn activate MAPKs at the TE/DY motif (Hamel et al., 2006). This hierarchical cascade ensures precise signal transmission, with MAPKKKs as initiators, MAPKKs as intermediates, and MAPKs as terminal effectors that phosphorylate substrates to regulate cellular responses (Rodriguez et al., 2010). MAPKs regulates growth, development, immunity, and defense in response to environmental and endogenous cues (Sinha et al., 2011; Manna et al., 2023). Evidence from multiple plant systems highlights MAPKs as important regulators of photosynthesis. In *Nicotiana attenuata* and *Arabidopsis*, MPK4 influences stomatal conductance, thylakoid organization, and PSII efficiency, thereby affecting carbon assimilation and energy distribution (Hettenhausen et al., 2012; Gawronski et al., 2014; Witoń et al., 2016, 2021). Moreover, *StMPK3*, *StMPK11*, and *PeMPK7*, modulate photosynthetic gene expression and stomatal dynamics under stress (Zhu et al., 2020, 2021; Gao et al., 2024). Furthermore, MPK4, MPK12, regulate CO₂-induced stomatal behaviour through Raf-like MAPKKK *HT1* (Hashimoto et al., 2016; Hiyama et al., 2017; K. Tõldsepp et al., 2018; Takahashi et al., 2022). MPK3 and MPK6 are rapidly activated during stress and immunity and often correlate with reduced photosynthetic activity (Su et al., 2018). Consistent with this, MPK3 overexpression has been shown to strongly reduce photosynthetic performance, whereas MPK6 exerts a modest positive effect, highlighting their distinct roles even under non-stressful conditions (Jonwal et al., 2023). However, the precise molecular mechanisms by which MAPKs influence photosynthesis remain poorly understood.

Here, we uncover a signaling framework that connects cytosolic MAPK activity to the photosynthetic capacity. We demonstrated that MPK3 modulates the activity of the Raf-like kinase ACTPK1, a cytosolic serine/threonine kinase previously implicated in nutrient signaling through its regulation of ammonium transporters (Beier et al., 2018). Notably, Arabidopsis homologues of ACTPK1 (STY8, STY17 and STY46) have been associated with chloroplast protein targeting and photosynthetic function (Lamberti et al., 2011), suggesting a broader role for this kinase family in organelle biogenesis. Building on these observations, our findings position ACTPK1 at the interface of MAPK signaling and chloroplast protein import. We show that MPK3 constrains ACTPK1 activity, thereby influencing the stability and targeting efficiency of the Rubisco small subunit precursor (RbcS). Our results further indicate that phosphorylation of the RbcS transit peptide operates as a reversible regulatory switch rather than a static modification, revealing a dynamic control mechanism that coordinates cytosolic signaling inputs with chloroplast protein import and photosynthetic output in rice

## Results

### MPK3 negatively regulates photosynthetic Carbon Assimilation

Overexpression of *MPK3* has been reported to adversely affects photosynthetic performance in rice (Jonwal et al., 2023). To dissect the molecular mechanism underlying this observation, we analyzed the photosynthesis and related parameters in two independent *MPK3* knockout lines (*mpk3 ko1* and *mpk3 ko2*) and two dexamethasone (DEX)-inducible *MPK3* overexpression lines (*MPK3-23* and *MPK3-24*) under ambient growth conditions (Banerjee et al., 2025; Singh et al., 2019). Photosynthetic pigments (chlorophyll a, chlorophyll b, and carotenoids) essential for light absorption and photoprotection were significantly higher in *mpk3 ko* and reduced in *MPK3 OE* lines compared to WT (Supplemental Fig. S1A-D). Elevated levels in knockout lines likely enhance light-harvesting efficiency and overall photosynthetic capacity, while reduced levels in overexpression lines may impair these processes. Consistently, net CO₂ assimilation rates (*A*) were significantly higher in *mpk3 ko* lines and lower in *MPK3 OE* lines than in WT (Fig. 1A). Under ambient CO₂ (400 µmol mol⁻¹), *A* reached 29.78 and 30.48 µmol m⁻² s⁻¹ in *mpk3 ko1* and *ko2* (18–21% higher than WT), whereas *MPK3 OE* lines showed only 17.75 and 17.67 µmol m⁻² s⁻¹ (∼29% lower than WT; Table 1).

**Fig. 1.**
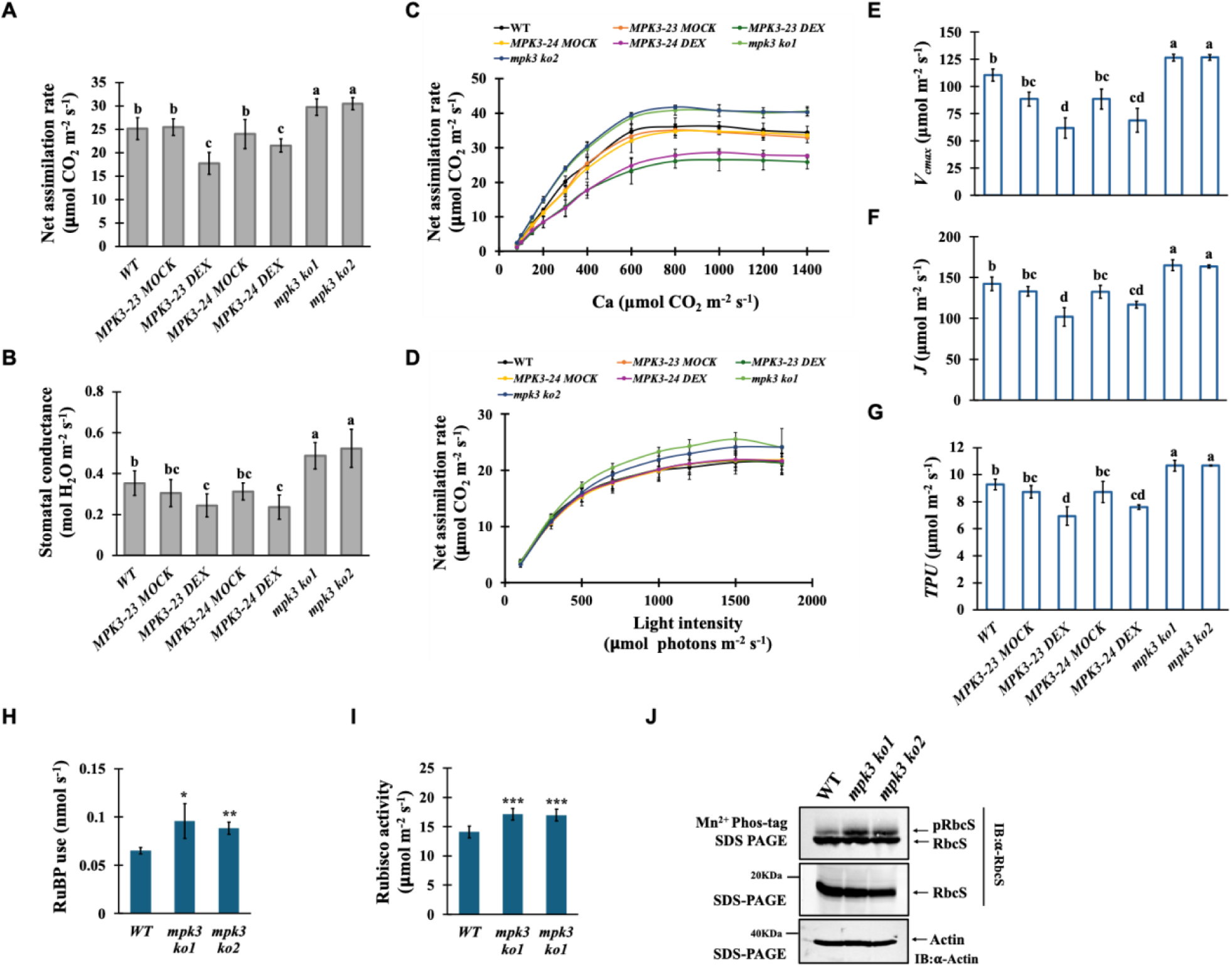
MPK3 negatively regulates photosynthesis. **A)** Bar graph showing net assimilation rate in WT, *MPK3 OE* (*MPK3-23*, *MPK3-24*) and *mpk3 ko* (*mpk3 ko1*, *mpk3 ko2*) plants at ambient CO_2_ (400 μmol mol^−1^). **B)** Bar graph showing Stomatal conductance to water in WT, *MPK3 OE* (*MPK3-23*, *MPK3-24*) and *mpk3 ko* (*mpk3 ko1*, *mpk3 ko2*) plants at ambient CO_2_ (400 μmol mol^−1^). **C)** Quantification of net photosynthesis per unit area of WT, *MPK3 OE* (*MPK3-23*, *MPK3-24*) and *mpk3 ko* (*mpk3 ko1*, *mpk3 ko2*) plants in response to varying CO_2_ concentration at 1500 μmol m^−2^ s^−1^ light intensity (*A/Ci* curve). Each value represents mean ± SD, where n = at least three data points from different plants. **D)** Quantification of net photosynthesis per unit area of WT, *MPK3 OE* (*MPK3-23*, *MPK3-24*) and *mpk3 ko* (*mpk3 ko1*, *mpk3 ko2*) plants in response to increasing PPFD at 400 μmol mol^−1^ CO_2_ concentration (*A/Q* curve). Each value represents mean ± SD, where n = at least three data points from different plants. **E)** Bar graph showing maximum rate of carboxylation (*Vcmax*) in WT, *MPK3 OE* (*MPK3-23*, *MPK3-24*) and *mpk3 ko* (*mpk3 ko1*, *mpk3 ko2*) plants. **F)** Bar graph showing electron transport rate (*J*) in WT, *MPK3 OE* (*MPK3-23*, *MPK3-24*) and *mpk3 ko* (*mpk3 ko1*, *mpk3 ko2*) plants. **G)** Bar graph showing triose phosphate utilisation (*TPU*) in WT, *MPK3 OE* (*MPK3-23*, *MPK3-24*) and *mpk3 ko* (*mpk3 ko1*, *mpk3 ko2*) plants. Data represented in D), E) and F) are deduced by fitting *A/Ci* curve using FvcB model. MOCK, dimethyl sulfoxide treated (DMSO) plants; DEX, dexamethasone treated plants. 1 µM DMSO and DEX sprayed on *MPK3 OE* leaves, 24 h prior to taking measurements. All the measurements were taken from flag leaf at heading stage using LiCOR6800, between 8:00 am to 11:00 am. Different letters indicate statistical significance according to one-way ANOVA followed by post-hoc Tukey HSD calculation at *P < 0.05*. **H)** Bar graph showing RuBP use in WT, *MPK3 OE* (*MPK3-23*, *MPK3-24*) and *mpk3 ko* (*mpk3 ko1*, *mpk3 ko2*) plants. **I)** Bar graph showing Rubisco activity in WT, *MPK3 OE* (*MPK3-23*, *MPK3-24*) and *mpk3 ko* (*mpk3 ko1*, *mpk3 ko2*) plants. Values represent the mean ± SD (*n* = 3 biological replicates). Asterix indicate statistical significance according to student ttest (*P*<0.05). **J)** Phos-tag gel western blots showing phosphorylated and non-phosphorylated RbcS content in WT and *mpk3 ko* lines. Blots were developed by α-RbcS antibody. Actin was used as endogenous control.

**Table 1:** CO_2_-saturated maximum photosynthesis (*A*_max_), maximum *in vivo* Rubisco activity (*V*_cmax_), maximum electron transport rate (*J*), and maximum rate of triose phosphate utilization (*TPU*) of WT, *MPK3 OE* and *mpk3 ko* lines through *A/C*_i_ curve analysis.

|  | *A<sub>mean</sub>* | *A<sub>max</sub>* | *V<sub>cmax</sub>* | *J* | *TPU* |
| --- | --- | --- | --- | --- | --- |
| **WT** | 25.17 ± 2.34 | 36.18 ± 1.37 | 110.2 ± 5.58 | 142.4 ± 8.23 | 9.28 ± 0.38 |
| **MPK3-23 MOCK** | 25.48 ± 1.8 | 35.16 ± 1.6 | 88.29 ± 6.348 | 133.16 ± 5.75 | 8.73 ± 0.45 |
| **MPK3-23 DEX** | 17.75 ± 2.34 | 26.53 ± 3.2 | 61.77 ± 9.32 | 101.89 ± 11.5 | 6.94 ± 0.688 |
| **MPK3-24 MOCK** | 24.02 ± 3.10 | 34.22 ± 0.87 | 88.368 ± 9.23 | 132.72 ± 7.9 | 8.73 ± .078 |
| **MPK3-24 DEX** | 17.67 ± 1.43 | 28.57 ± 0.51 | 68.79 ± 10.83 | 116.92 ± 4.18 | 7.6 ± 0.17 |
| ***mpk3 ko1*** | 29.78 ± 1.75 | 40.83 ± 1.41 | 126.3 ± 3.2 | 165.12 ± 6.58 | 10.66 ± 0.4 |
| ***mpk3 ko2*** | 30.48 ± 1.28 | 41.8 ± 0.4 | 126.56 ± 2.54 | 163.65 ± 1.9 | 10.66 ± 0.02 |

Photosynthetic efficiency is determined by CO₂ diffusion into the chloroplast stroma (stomatal conductance, gs) and subsequent carboxylation (Evans, 2021). To explore the basis for the observed differences in *A*, we measured *gs* and intercellular CO₂ concentration (*Ci*). Notably, *mpk3 ko* plants with higher *A* exhibited elevated *gs* whereas *MPK3 OE* plants showed reduced *gs* compared to WT. However, *Cᵢ* values remained comparable across genotypes (Fig. 1B and Supplemental Fig. S1E), indicating that the altered photosynthetic rates stem primarily from differences in carboxylation efficiency rather than CO₂ availability.

Since *mpk3 ko* plants showed higher *A* and *gs*, but unchanged *Ci*, we used *A/Ci* and *A/Q* curve analyses to distinguish the effects of MPK3 on CO₂ fixation versus light-dependent processes (Farquhar et al., 1980; Long & Bernacchi, 2003). With increasing CO_2_, *mpk3 ko* plants exhibited consistently higher *A* compared to WT, with a CO₂-saturated maximum assimilation rate (*Amax*) of 41.31 ± 0.68 µmol CO₂ m⁻² s⁻¹ (∼14% higher than WT), while *MPK3 OE* lines showed a reduced *Amax* of 27.55 ± 1.45 µmol CO₂ m⁻² s⁻¹ (∼31% lower than WT) (Fig. 1C and Table 1). In contrast, light-response curves were similar among genotypes, with comparable light-saturated assimilation rates (Fig. 1D), indicating that MPK3 primarily modulates CO₂-responsive processes, such as Calvin cycle efficiency, rather than light-dependent reactions.

Consistent with the preferential effect of MPK3 on CO₂-responsive photosynthesis, Farquhar-von Caemmerer-Berry (FvCB) modelling of the *A/Ci* curves revealed that *Vcmax*, *J*, and *TPU* were significantly enhanced in *mpk3 ko* plants but markedly reduced in *MPK3 OE* lines relative to WT (Fig. 1E, F & G), indicating that MPK3 negatively regulates the biochemical capacity for CO₂ fixation and regeneration in the Calvin cycle. These results establish *MPK3* as a negative regulator of photosynthetic efficiency, as knockout lines exhibited enhanced pigment accumulation, stomatal conductance, and carboxylation efficiency, boosting photosynthesis, while overexpression lines showing the opposite trend.

### MPK3 Regulates Rubisco Activity via RbcS Phosphorylation

Carboxylation efficiency is primarily governed by Rubisco activity and the availability of its substrate RuBP (Farquhar et al., 1980). Supporting the higher *Vcmax* in *mpk3 ko* plants, both RuBP utilisation and Rubisco activity were significantly increased compared with WT (Fig. 1H, I). The observation indicate that MPK3 adversely affect Rubisco function and RuBP regeneration, thereby limits carboxylation efficiency. The fact that Rubisco activity depends on the assembly of the nuclear-encoded small subunit (RbcS) with the chloroplast-encoded large subunit (RbcL), and that MPK3 localises predominantly to the nucleus and cytosol (Supplemental Fig. S2A, B), we hypothesised that MPK3 regulates Rubisco function via transcriptional or post-translational control of RbcS rather than through direct effects on chloroplast-resident RbcL.

Accordingly, to test whether MPK3 regulates RbcS post-translationally, we examined RbcS phosphorylation by Phos-tag analysis and detected an upshift of RbcS band indicative of phosphorylation in both WT and *mpk3 ko* plants. Notably, the phosphorylated RbcS band was consistently more intense in *mpk3 ko* plants compared to WT, suggesting enhanced RbcS phosphorylation in the absence of *MPK3* (Fig. 1J). This higher RbcS phosphorylation in *mpk3 ko* is positively correlated with higher Rubisco activity and carboxylation rates.

These findings imply that MPK3 does not directly phosphorylate RbcS but rather suppresses the activity of another kinase responsible for this modification. In *mpk3 ko* plants, where MPK3 is absent, this another “missing” kinase remains active, leading to increased RbcS phosphorylation. Given the functional overlap between MPK3 and MPK6, we tested whether MPK6 could act as this “missing” kinase. However, yeast two-hybrid (Y2H) and *in vitro* kinase assays revealed no interaction or phosphorylation of RbcS by either MPK3 or MPK6 (Supplemental Fig. S2C & D), ruling out MPK6 as the “missing” kinase. These findings strongly suggest that MPK3 indirectly modulates Rubisco activity by regulating RbcS phosphorylation via an unidentified kinase.

### ACTPK1 may act as a Candidate “missing” Kinase in MPK3-RbcS module

To identify the kinase responsible for elevated RbcS phosphorylation in *mpk3 ko* plants, we conducted a literature survey focusing on known kinases that modify RbcS or its precursor. In *Arabidopsis*, the RbcS transit peptide is phosphorylated by the cytosolic STY kinases, STY8, STY17 and STY46, which regulate chloroplast import and thereby influence Rubisco assembly and photosynthetic performance (Lamberti et al., 2011). A homology search using AtSTY46 identified a rice ACT-domain–containing kinase, ACTPK1 (LOC_Os02g02780) (Supplemental Fig. S3A), which clusters within the Group-C Raf-like MAPKKKs and shows strong sequence similarity to STY8, STY17 and STY46 (Supplemental Fig. S3B, C) (Beier et al 2018; Rao et al., 2010). Notably, ACTPK1 contains multiple canonical S/TP motifs corresponding to MAPK phosphorylation sites (Supplemental Fig. S3D), suggesting potential regulation by MPK3. Given these structural and sequence similarities, ACTPK1 emerges as a strong candidate for the kinase suppressed by MPK3 in regulating RbcS phosphorylation.

### MPK3 physically interacts with ACTPK1

To test whether ACTPK1 is regulated by MPK3, we first examined their physical interaction, as MAPKs typically regulate downstream targets through direct binding and phosphorylation (Manna et al., 2023). We initially assessed the interaction between ACTPK1 and MPK3 in yeast cell (*Saccharomyces cerevisiae*) using Y2H assay. ACTPK1 fused to the GAL4 DNA-binding domain (BD) and MPK3 fused to the GAL4 activation domain (AD), were co-transformed into the Y2H Gold strain in different combinations. Y2H assay showed that ACTPK1 physically interacted with MPK3 (Fig. 2A), with the p53–T7 pair and empty-vector combinations serving as positive and negative controls, respectively.

**Fig. 2.**
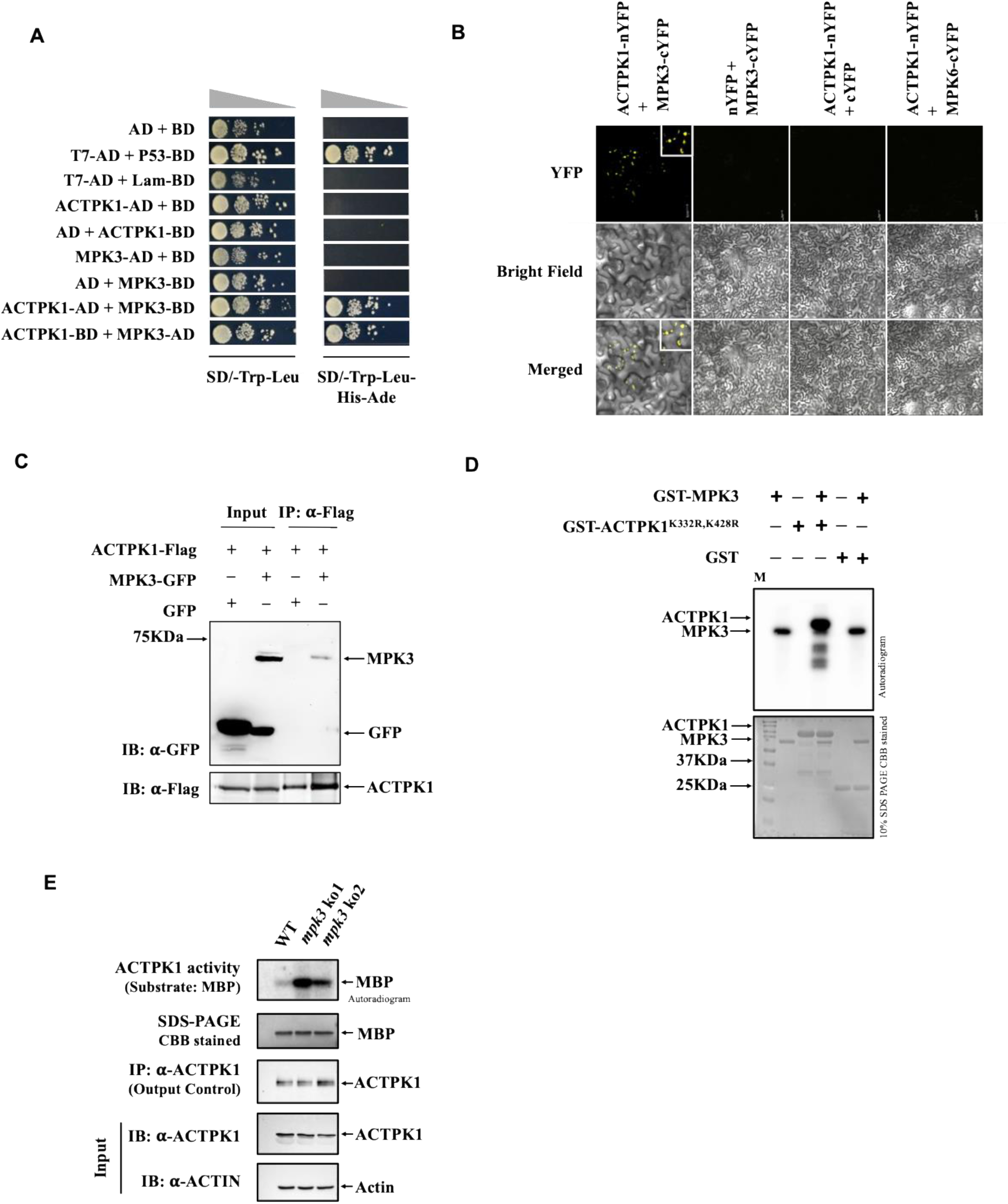
MPK3 physically interacts with and phosphorylates ACTPK1 to inhibit its activity. **A)** MPK3 interacts with ACTPK1 in yeast cells. Yeast cells were cultured on SD/-Trp-Leu and SD/-Trp-Leu-His-Ade media. AD, GAL4 activation domain; BD, GAL4 DNA-binding domain; SD, synthetic defined. **B)** BiFC assays showing that MPK3 interacts with ACTPK1 in *N. benthamiana*. ACTPK1-nYFP was co-transformed with MPK3-cYFP in fully expanded leaves of *N. benthamiana*. The ACTPK1-nYFP/MPK6-cYFP and Empty-nYFP/MPK3-cYFP were used as negative controls. Inset on top right is the enlarged image of signal. Scale Bar, 25 *µm*. **C)** Interaction between MPK3 and ACTPK1 in the Co-IP assays. α-Flag beads were used to immunoprecipitate ACTPK1 protein. Gel blots were probed with α-GFP antibody. **D)** *In vitro* kinase assay showing phosphorylation of kinase dead GST-ACTPK1^K332R,K428R^ by GST-MPK3. CBB (Coomassie Brilliant Blue) stained 10% SDS page gel shows loaded proteins. **E)** IP-Kinase showing ACTPK1 immunoprecipitated from *mpk3 ko* enhances MBP phosphorylation. ACTPK1 was immunoprecipitated using α-ACTPK1 antibody.

The interaction was confirmed *in planta* by bimolecular fluorescence complementation (BiFC) assay in *Nicotiana benthamiana.* ACTPK1 was fused to the N-terminal half of enhanced yellow fluorescent protein (nYFP), while MPK3 was fused to the C-terminal half (cYFP). Strong YFP fluorescence was detected in *N. benthamiana* cells co-expressing ACTPK1–nYFP and MPK3–cYFP, but not in control combinations, including ACTPK1 with MPK6 (Fig. 2B). The association was further validated by co-immunoprecipitation of ACTPK1–Flag with MPK3–GFP from *N. benthamiana* extracts, whereas no signal was detected in control samples (Fig. 2C). These results establish a specific physical interaction between MPK3 and ACTPK1, prompting us to examine whether MPK3 functionally regulates ACTPK1 by phosphorylation.

### MPK3 phosphorylates ACTPK1 and supress its kinase activity

As a putative Raf like MAPKKK, ACTPK1 was predicted to exhibit kinase activity. To test this possibility, the coding sequence of ACTPK1 was fused with glutathione S-transferase (GST) for protein purification. The resultant purified protein was subjected to *in vitro* kinase assay with myelin basic protein (MBP), a widely used universal kinase substrate (Eichberg and Iyer, 1996). Briefly, GST-ACTPK1 was incubated with MBP in kinase assay buffer supplemented with λ^32^P-ATP for 30 min at 30°C and the reaction products were resolved by SDS-PAGE. Autoradiography revealed strong phosphorylation signals corresponding to both ACTPK1 and MBP, indicating that ACTPK1 possesses both auto- and trans-phosphorylation activities (Supplemental Fig. S4A).

To test whether ACTPK1 is a direct substrate of MPK3, we generated catalytically inactive variants of ACTPK1 to eliminate background phosphorylation arising from its intrinsic kinase activity. To generate a catalytically inactive ACTPK1 variant, a conserved lysine in the predicted catalytic subdomain II (K332), identified by sequence alignment with homologues (Supplemental Fig. S4B), was mutated to arginine (ACTPK1^K332R^) by site-directed mutagenesis (Supplemental Fig. S4C). This mutation abolished MBP trans-phosphorylation while retaining ACTPK1 autophosphorylation (Supplemental Fig. S4A), indicating that K332 is required for trans-phosphorylation but not for autophosphorylation. To further eliminate residual autophosphorylation, a second conserved lysine in subdomain VI (K428) was mutated in the K332R background (ACTPK1^K332R,K428R^) (Supplemental Fig. S4D), which abolished both auto- and trans-phosphorylation (Supplemental Fig. S4A), generating a fully inactive ACTPK1 variant.

Using this kinase-dead ACTPK1^K332R,K428R^ protein as substrate for MPK3, an in-solution kinase assays demonstrated that MPK3 directly phosphorylates ACTPK1 (Fig. 2D), whereas GST alone was not phosphorylated, confirming the specificity of the reaction. In *Medicago sativa*, OMTK1 (a MAPKKK) was shown to directly phosphorylate and activate the MMK3 (a MAPK), bypassing the canonical MAPKK step (Nakagami et al., 2004). This represents a rare exception to the typical three-tier MAPK cascade and highlights alternative signalling modes in plants. Since the evidence of reverse signalling in MAPK modules is rare, we tested whether ACTPK1 could phosphorylate MPK3. However, ACTPK1 failed to phosphorylate a kinase-dead MPK3 variant (MPK3ᴷ⁶⁵ᴿ) (Supplemental Fig. S4E), indicating that phosphorylation occurs unidirectionally from MPK3 to ACTPK1.

To assess the functional consequence of MPK3-mediated phosphorylation of ACTPK1, we immunoprecipitated ACTPK1 from WT and *mpk3 ko* plants and measured its kinase activity using MBP as substrate. ACTPK1 immunocomplexes from *mpk3 ko* plants displayed substantially higher kinase activity than those from WT (Fig. 2E), demonstrating that MPK3 negatively regulates ACTPK1 kinase activity, likely through direct phosphorylation-mediated inhibition.

### ACTPK1 interacts with and phosphorylates RbcS at its signal peptide

To establish ACTPK1 as the kinase responsible for RbcS phosphorylation in our pathway, we analysed *RbcS3*, the most abundant *RbcS* isoform in rice (∼50% of the total RbcS pool; Suzuki et al., 2007). Since ACTPK1 is a cytosolic kinase, we reasoned that its interaction with RbcS would occur in the cytoplasm prior to plastid import and therefore used the precursor form of RbcS for all interaction assays. Y2H analysis revealed a direct interaction between ACTPK1 and precursor RbcS (Fig. 3A). This interaction was further confirmed *in planta* by BiFC assay in *N. benthamiana*, where a strong YFP signal was detected in the cells co-expressing ACTPK1–nYFP and RbcS–cYFP (Fig. 3B), demonstrating that ACTPK1 physically associates with RbcS *in vivo*.

**Fig. 3.**
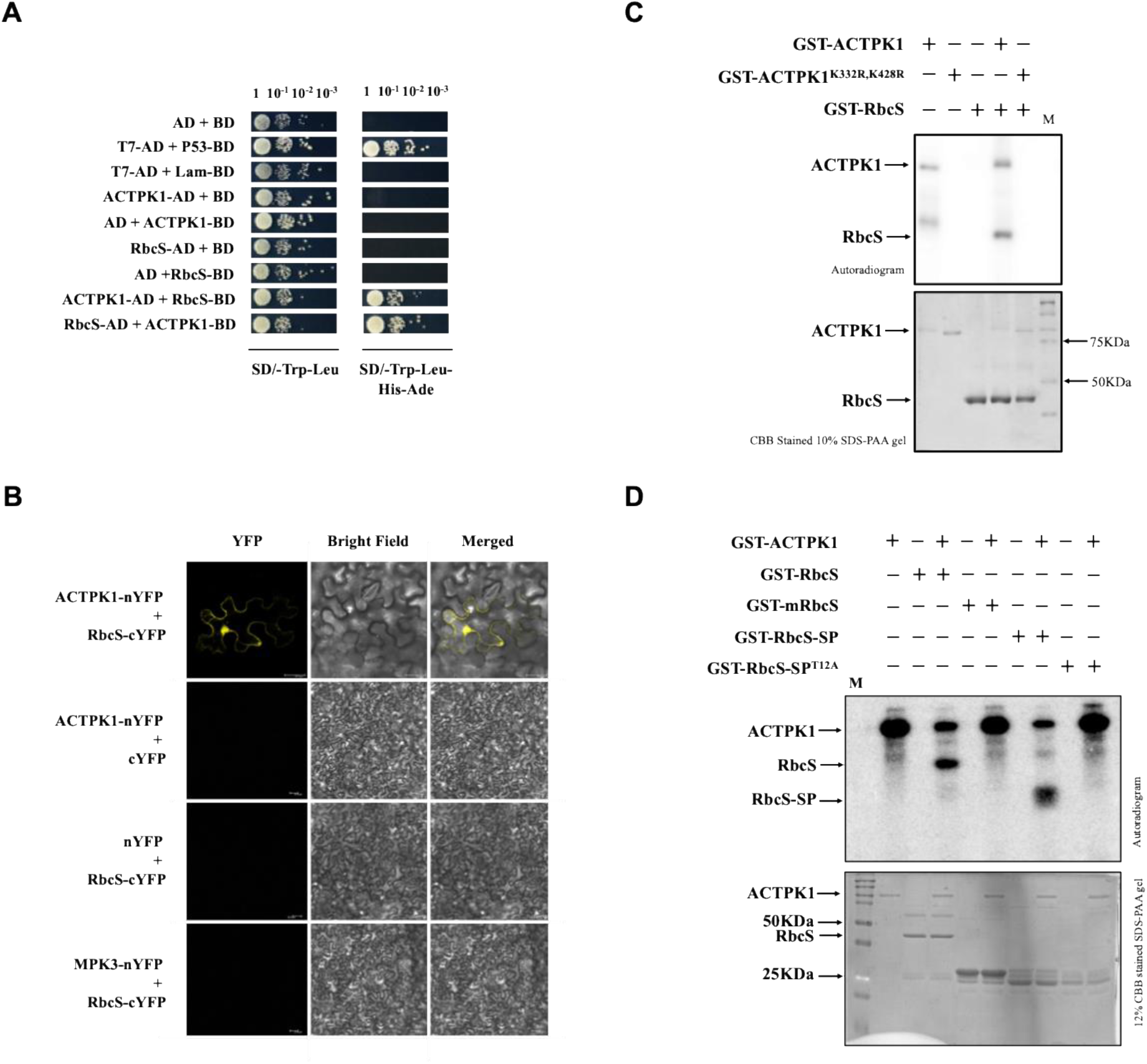
ACTPK1 physically interacts with and phosphorylates RbcS within signal peptide. **A)** ACTPK1 interacts with RbcS in yeast cells. Yeast cells were cultured on SD/-Trp-Leu and SD/-Trp-Leu-His-Ade media. AD, GAL4 activation domain; BD, GAL4 DNA-binding domain; SD, synthetic defined. **B)** BiFC assays showing that ACTPK1 interacts with RbcS in *N. benthamiana* mesophyll cell. ACTPK1-nYFP was co-transformed with RbcS-cYFP in fully expanded leaves of *N. benthamiana*. The ACTPK1-nYFP/cYFP, nYFP/RbcS-cYFP and MPK3-nYFP/RbcS-cYFP were used as negative controls. Scale Bar, 25 *µ*m. **C)** *In vitro* kinase assay showing phosphorylation of kinase dead GST-RbcS by GST-ACTPK1. CBB stained 10% SDS PAA gel shows loaded proteins. **D)** *In vitro* kinase assay showing ACTPK1 specifically phosphorylates Thr12 residue of RbcS. CBB stained 12% SDS PAA gel shows loaded proteins.

We next tested whether this interaction results in phosphorylation of RbcS. *In vitro* kinase assays using bacterially expressed GST-ACTPK1 and GST-RbcS showed robust phosphorylation, demonstrating that ACTPK1 directly phosphorylates RbcS (Fig. 3C). To pinpoint the region of phosphorylation, we cloned and expressed precursor RbcS (containing the signal peptide) and its mature form lacking the signal peptide (mRbcS). *In vitro* kinase assay revealed that phosphorylation occurred exclusively on the precursor but was abolished in mRbcS, indicating that the phosphorylation site resides within the signal peptide (Supplemental Fig. S5A,B). *In silico* analysis using NetPhos3.1 identified three high-confidence candidate phosphosites within the signal peptide (Thr12, Ser31 and Ser34; Supplemental Fig. S5C). Alanine-substitution mutants (RbcS^S31A^, RbcS^S34A^ and RbcS^T12A^) were generated in precursor RbcS form and subsequently tested in kinase assays (Supplemental Fig. S5D). Phosphorylation was retained in RbcS^S31A^ and RbcS^S34A^ but completely abolished in RbcS^T12A^ (Fig. 3D; Supplemental Fig. S5E).

Together, these findings demonstrate that ACTPK1 directly phosphorylates RbcS at Thr12 within its signal peptide. This modification likely act as a regulatory checkpoint in the cytoplasm, modulating RbcS import into chloroplasts and thereby fine-tuning Rubisco assembly and function.

### Phosphorylation by ACTPK1 regulates RbcS stability and chloroplast targeting competence

Since ACTPK1 phosphorylates RbcS at Thr12 within its chloroplast transit peptide, a region essential for TOC/TIC-mediated targeting, we tested whether modification of this residue affects RbcS chloroplast import. To this end, phospho-dead (RbcS^T12A^) and phospho-mimetic (RbcS^T12D^) variants of the RbcS precursor were generated by site directed mutagenesis using overlapping PCR (Fig. 4A). The mutant versions along with Wild-type were fused to C-terminal GFP and transiently expressed in rice protoplasts derived from WT, *mpk3 ko*, and *actpk1 ko* backgrounds. Wild-type RbcS–GFP predominantly localized to chloroplasts, whereas both the phospho-dead (RbcS^T12A^) and phospho-mimetic (RbcS^T12D^) variants displayed markedly reduced chloroplast accumulation and instead localized primarily to the cytosol and perichloroplast regions (Fig. 4B). Notably, the phospho-dead RbcS^T12A^ variant accumulated in distinct cytosolic puncta, suggesting aggregation of precursor protein, whereas the phospho-mimetic RbcS^T12D^ variant remained diffusely distributed in the cytosol but still failed to efficiently localize to chloroplasts. These observations suggest that phosphorylation of Thr12 may help maintain the RbcS precursor in a soluble, import-competent state, while subsequent dephosphorylation may be required for productive chloroplast import. The impaired targeting of both phospho-dead and phospho-mimetic variants therefore indicates that efficient chloroplast import does not depend on a fixed phosphorylation state but requires regulated phosphorylation dynamics. Supporting this interpretation, pharmacological inhibition of protein dephosphorylation by okadaic acid results in reduced chloroplast accumulation of wild-type RbcS–GFP, phenocopying the localization defect observed for the phospho-mimetic variant (Fig. 4C). Moreover, the ability of the phospho-dead and phospho-mimetic mutant to partly localize to chloroplasts indicates that phosphorylation is not strictly required for import. However, its reduced efficiency compared to the wild-type RbcS suggests that dynamic phosphorylation–dephosphorylation, rather than a constitutive state, optimizes chloroplast targeting. Similar localization pattern was observed in *mpk3 ko* protoplasts (Supplemental Fig. S6A).

**Fig. 4.**
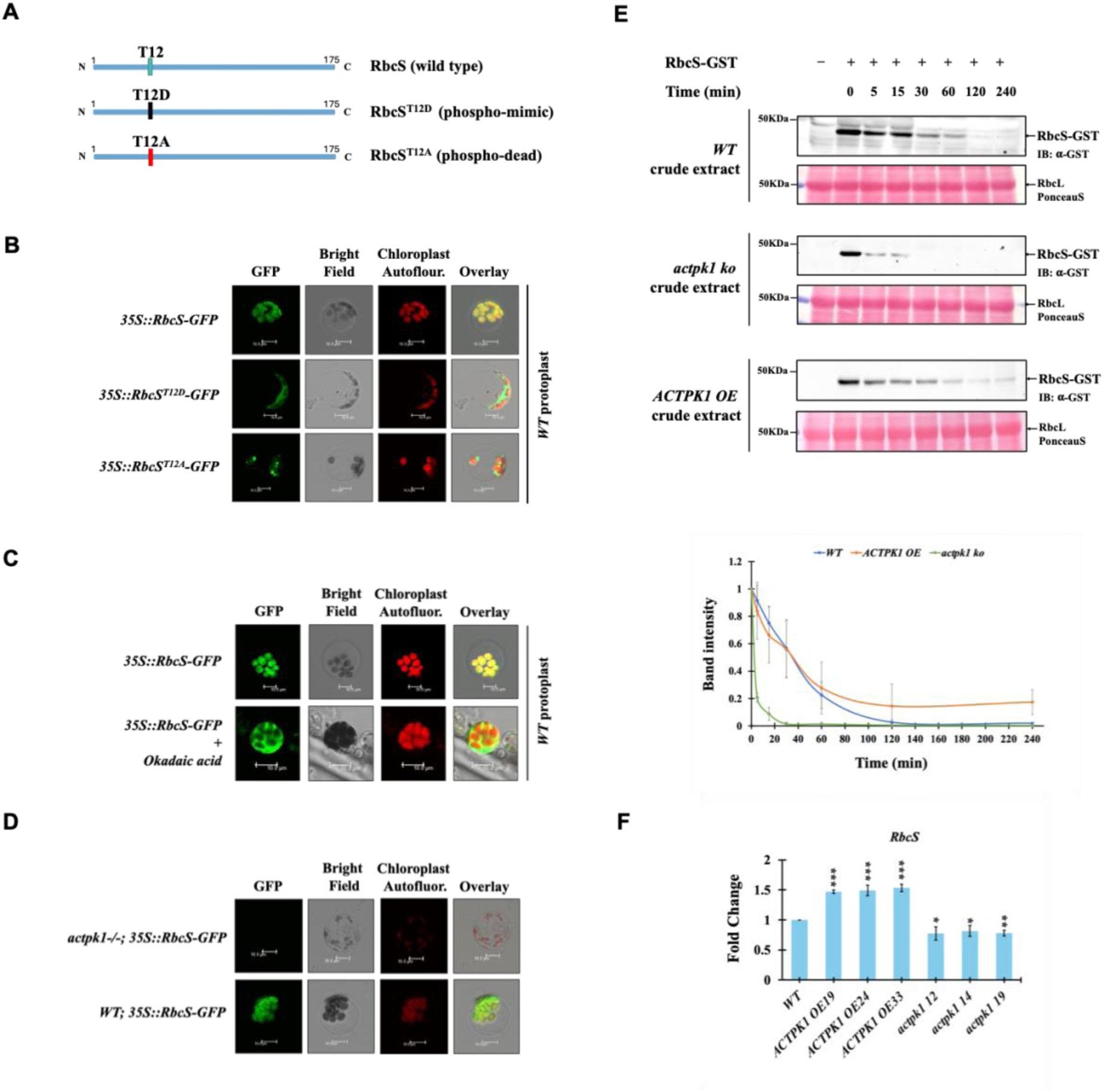
ACTPK1 regulates stability and import of RbcS into the chloroplast. **A)** Pictorial representation of RbcS, RbcS^T12D^ (phosphor-mimic), RbcS^T12A^ (phosphor-dead) mutant constructs designed through site-directed mutagenesis. **B)** Localization of RbcS, RbcS^T12D^, RbcS^T12A^ in protoplast isolated from WT rice seedlings. Scale Bar, 10 *µ*m. **C)** Localization of RbcS in protoplast of WT rice seedlings treated with or without okadaic acid. 50nM okadaic acid for 30 min was used for treating WT protoplasts transfected with RbcS-GFP. Scale Bar, 10 *µ*m. **D)** Localization of RbcS in the protoplasts of stable transgenic lines of WT and *actpk1 ko* overexpressing *RbcS* under the control of 35SCaMV promoter. Scale Bar, 10 *µ*m. **E)** Western blots of cell-free degradation assays showing degradation of GST-RbcS when incubated with crude extract of WT (upper panel), *actpk1 ko* (middle panel) and *ACTPK1 OE* (lower panel). Ponceau stained (PonceauS) blot shows RbcL protein. Experiment was repeat thrice. **E)** Line graph depicting the band intensities as shown in (E). Band intensities were quantified using ImageJ software. **F)** Bar graph showing the RbcS transcript abundance in WT, *ACTPK1 OE* (*ACTPK1 OE19, ACTPK1 OE24 & ACTPK1 OE33*) and *actpk1 ko* (*actpk1 ko12, actpk1 ko14 & actpk1 ko19*). Actin7 and Ubq5 were used as internal control. Values represent the mean ± SD (*n* = 3 biological replicates). Asterix indicate statistical significance according to student ttest (*P*<0.05).

Strikingly, no detectable GFP signal from RbcS–GFP was observed in *actpk1 ko* protoplasts, despite multiple independent transfection attempts (Supplemental Fig. S6B). In line with this observation, stable transgenic lines expressing RbcS–GFP showed robust chloroplast localization in the WT background but no detectable signal in the *actpk1 ko* background (Fig. 4D). Treatment with the proteasome inhibitor MG132 did not restore GFP accumulation, indicating that proteasome-dependent degradation alone does not fully account for the loss of RbcS–GFP signal (Supplemental Fig. S6B).

To directly examine the RbcS stability, we performed a cell-free protein degradation assay by incubating bacterially expressed RbcS protein with crude extracts from WT, *ACTPK1 OE* and *actpk1 ko* plants. RbcS was rapidly destabilized in extracts derived from *actpk1 ko* plants, whereas it remained comparatively stable in WT and *ACTPK1 OE* extracts (Fig. 4E). These results indicate that ACTPK1 activity contributes to maintaining RbcS protein stability.

To further substantiate these observations, quantitative RT-PCR analysis revealed reduced RbcS transcript levels in *actpk1 ko* plants compared with WT and *ACTPK1 OE* lines (Fig. 4F), suggesting that ACTPK1 influences RbcS accumulation at multiple regulatory levels. Together, these findings demonstrate that ACTPK1-dependent phosphorylation is required to maintain RbcS stability and chloroplast targeting competence, thereby supporting efficient precursor accumulation and import.

### ACTPK1 positively regulates photosynthesis

To further validate this pathway and its impact on photosynthesis, we generated *ACTPK1* overexpression (*OE*) and knockout (*ko*) lines, along with *MPK3-ACTPK1* double knockout (*mpk3actpk1 ko*) *in the* japonica rice cultivar Taipae309. Three independent *OE* lines (*ACTPK1 OE19*, *ACTPK1 OE24* and *ACTPK1 OE33*) with the highest transcript accumulation were selected for detailed analysis (Supplemental Fig. S7A, B & C). *actpk1 ko* lines were generated via CRISPR/Cas9-mediated genome editing technology and three independent *actpk1* mutant alleles, *actpk1 ko12 (*deletion of four nucleotides*), actpk1 ko14* (single nucleotide insertion) and *actpk1 ko19* (single nucleotide insertion), were selected for detailed analysis (Supplemental Fig. S7D & E). The double knockout was generated by mutating ACTPK1 in *mpk3 ko* background using CRISPR-Cas9 genome editing tool. A mutant line harbouring a single-nucleotide insertion in the ACTPK1 coding sequence was selected for further analysis (Supplemental Fig. S7F).

To study the impact of ACTPK1 on photosynthesis we measured gas-exchange parameters in the flag leaf of *ACTPK1 OE* and *actpk1 ko* lines at heading stage using LI-COR 6800 system, targeting a 2 cm² area from 9:00 am to 11:00 am. At ambient CO_2_ concentration, the mean *A* of *actpk1 ko* plants were significantly lower that WT and *ACTPK1 OE* lines, with *actpk1 ko* showing 12.973 ± 0.99 μmol m⁻² s⁻¹, a 36–45% decrease compared to WT and *ACTPK1 OE* plants (Fig. 5A & Table 2). In contrast, mean *A* in *ACTPK1 OE* lines were comparable to WT, indicating that overexpression does not disturb the kinase’s functional balance.

**Fig. 5.**
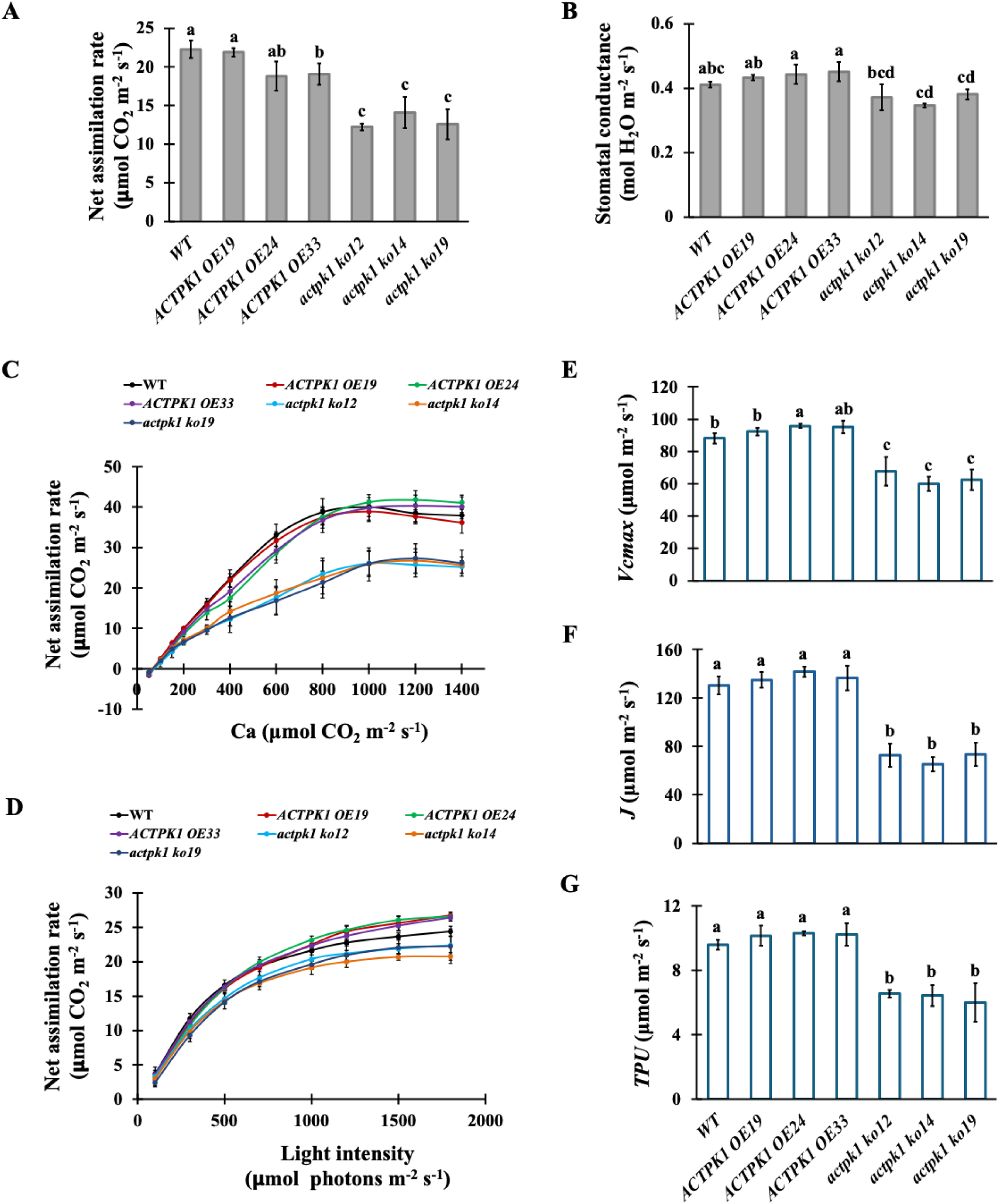
ACTPK1 positively regulates photosynthesis in rice. **A)** Bar graph showing net assimilation rate in WT, *ACTPK1 OE* (*ACTPK1 OE19, ACTPK1 OE24 & ACTPK1 OE33*) and *actpk1 ko* (*actpk1 ko12, actpk1 ko14 & actpk1 ko19*) plants at ambient CO_2_ (400 μmol mol^−1^). **B)** Bar graph showing Stomatal conductance to water in WT, *ACTPK1 OE* (*ACTPK1 OE19, ACTPK1 OE24 & ACTPK1 OE33*) and *actpk1 ko* (*actpk1 ko12, actpk1 ko14 & actpk1 ko19*) plants at ambient CO_2_ (400 μmol mol^−1^). **C)** Quantification of net photosynthesis per unit area of WT, *ACTPK1 OE* (*ACTPK1 OE19, ACTPK1 OE24 & ACTPK1 OE33*) and *actpk1 ko* (*actpk1 ko12, actpk1 ko14 & actpk1 ko19*) plants in response to varying CO_2_ concentration at 1500 μmol m^−2^ s^−1^ light intensity (*A/Ci* curve). Each value represents mean ± SD (n = at least three biological replicate). **D)** Quantification of net photosynthesis per unit area of WT, *ACTPK1 OE* (*ACTPK1 OE19, ACTPK1 OE24 & ACTPK1 OE33*) and *actpk1 ko* (*actpk1 ko12, actpk1 ko14 & actpk1 ko19*) plants in response to increasing PPFD at 400 μmol mol^−1^ CO_2_ concentration (*A/Q* curve). Each value represents mean ± SD (n = at least three biological replicate). **E)** Bar graph showing maximum rate of carboxylation (*Vcmax*) in WT, *ACTPK1 OE* (*ACTPK1 OE19, ACTPK1 OE24 & ACTPK1 OE33*) and *actpk1 ko* (*actpk1 ko12, actpk1 ko14 & actpk1 ko19*) plants. **F)** Bar graph showing electron transport rate (*J*) in WT, *ACTPK1 OE* (*ACTPK1 OE19, ACTPK1 OE24 & ACTPK1 OE33*) and *actpk1 ko* (*actpk1 ko12, actpk1 ko14 & actpk1 ko19*) plants. **G)** Bar graph showing triose phosphate utilisation (*TPU*) in WT, *ACTPK1 OE* (*ACTPK1 OE19, ACTPK1 OE24 & ACTPK1 OE33*) and *actpk1 ko* (*actpk1 ko12, actpk1 ko14 & actpk1 ko19*) plants. Data represented in E), F) and G) are deduced by fitting *A/Ci* curve using FvcB model. All the measurements were taken from flag leaf at heading stage using LiCOR6800, between 8:00 am to 11:00 am. Different letters indicate statistical significance according to one-way ANOVA followed by post-hoc Tukey HSD calculation at *P < 0.05*.

**Table 2:** CO_2_-saturated maximum photosynthesis (*A*_msax_), maximum *in vivo* Rubisco activity (*V*_cmax_), maximum electron transport rate (*J*), and maximum rate of triose phosphate utilization (*TPU*) of WT, *ACTPK1 OE* and *actpk1 ko* lines through *A/C*_i_ curve analysis.

|  | *A<sub>mean</sub>* | *A<sub>max</sub>* | *V<sub>cmax</sub>* | *J* | *TPU* |
| --- | --- | --- | --- | --- | --- |
| **WT** | 22.27 ± 1.12 | 38.8 ± 1.08 | 88.18 ± 3.14 | 130.36 ± 7.36 | 9.5 ± 0.31 |
| **ACTPK1 OE19** | 21.89 ± 0.55 | 41.34 ± 0.37 | 88.25 ± 2.48 | 124.79 ± 6.37 | 9.13 ± 0.61 |
| **ACTPK1 OE24** | 18.42 ± 1.17 | 39.98 ± 0.26 | 105.77 ± 1.29 | 141.49 ± 4.39 | 10.28 ± 0.12 |
| **ACTPK1 OE33** | 19.08 ± 0.56 | 37.5 ± 1.36 | 95.05 ± 3.88 | 133.93 ± 12.95 | 10.3 ± 0.96 |
| ***actpk1 ko12*** | 12.22 ± 0.42 | 25.65 ± 0.44 | 67.6 ± 8.85 | 72.58 ± 9.65 | 6.53 ± 0.23 |
| ***actpk1 ko14*** | 14.10 ± 2.03 | 26.03 ± 0.54 | 59.89 ± 4.42 | 65.23 ± 5.89 | 6.42 ± 0.65 |
| ***actpk1 ko19*** | 12.59 ± 1.93 | 26.48 ± 0.7 | 62.38 ± 6.36 | 73.32 ± 9.63 | 5.98 ± 1.19 |

Photosynthesis is positively corelated with stomatal conductance. Stomatal conductance mirrored these patterns, with *actpk1 ko* lines showing lower values supporting the reduced photosynthetic capacity (Fig. 5B). *Ci* remained unchanged among genotypes, ruling out CO₂ availability as the cause of reduced photosynthesis in *actpk1 ko* plants. (Supplemental Fig. 8). Moreover, *A/Ci* response curves revealed a pronounced reduction in net photosynthetic rate in *actpk1 ko* lines compared to WT and *ACTPK1 OE* lines across all tested CO₂ concentrations (Fig. 5C). Interestingly, *ACTPK1 OE* lines exhibited photosynthetic rates comparable to WT across all tested CO₂ concentrations. *A/Q* curves further showed that *actpk1 ko* lines had diminished light-saturated photosynthesis, while *ACTPK1 OE* lines sustained slightly higher rates relative to WT (Fig. 5D).

Modelling of the *A/Ci* curves provided further insight into the biochemical basis of this decline. The CO₂-saturated photosynthetic capacity (*Aₘₐₓ)* was reduced by ∼31–37% in *actpk1 ko* plants (26.2 ± 0.5 μmol m⁻² s⁻¹) compared with WT (39.3 ± 0.8 μmol m⁻² s⁻¹) and *OE* lines (39.9 ± 1.5 μmol m⁻² s⁻¹) (Table 2). Moreover, *actpk1 ko* lines displayed significantly reduced values for *Vcmax*, *J*, and *TPU* (Fig. 5E, F & G). These reductions strongly suggest impaired Rubisco activity and compromised electron transport in *actpk1 ko* plants. Collectively, these findings demonstrate that the absence of ACTPK1 leads to a substantial loss in Rubisco-dependent photosynthetic capacity.

### MPK3-ACTPK1 operate in a linear signaling cascade to regulate photosynthesis

Next, we analysed the photosynthetic phenotypes of *mpk3*, *actpk1*, and *mpk3actpk1* mutants. *A/Ci* response curves revealed a marked increase in net CO₂ assimilation in *mpk3 ko* plants, with a steep rise and elevated plateau, whereas *actpk1 ko* lines displayed strongly reduced photosynthetic rates relative to WT, across a range of intercellular CO₂ concentrations (Fig. 6A). Crucially, the *mpk3actpk1* double mutant mirrors the *actpk1 ko* phenotype, with low assimilation rate, rather than the enhanced *mpk3* ko mutant. This suppression in the *mpk3actpk1* double mutant indicates that ACTPK1 works downstream of MPK3 as in absence of ACTPK1, knocking out MPK3 cannot rescue the pathway. A *s*imilar trend was observed for light response (*A/Q*) curves where *mpk3 ko* plants maintained slightly elevated photosynthetic rates under increasing irradiance, while *actpk1 ko* and *mpk3actpk1 ko* lines exhibited impaired light-saturated photosynthesis (Fig. 6B).

**Fig. 6.**
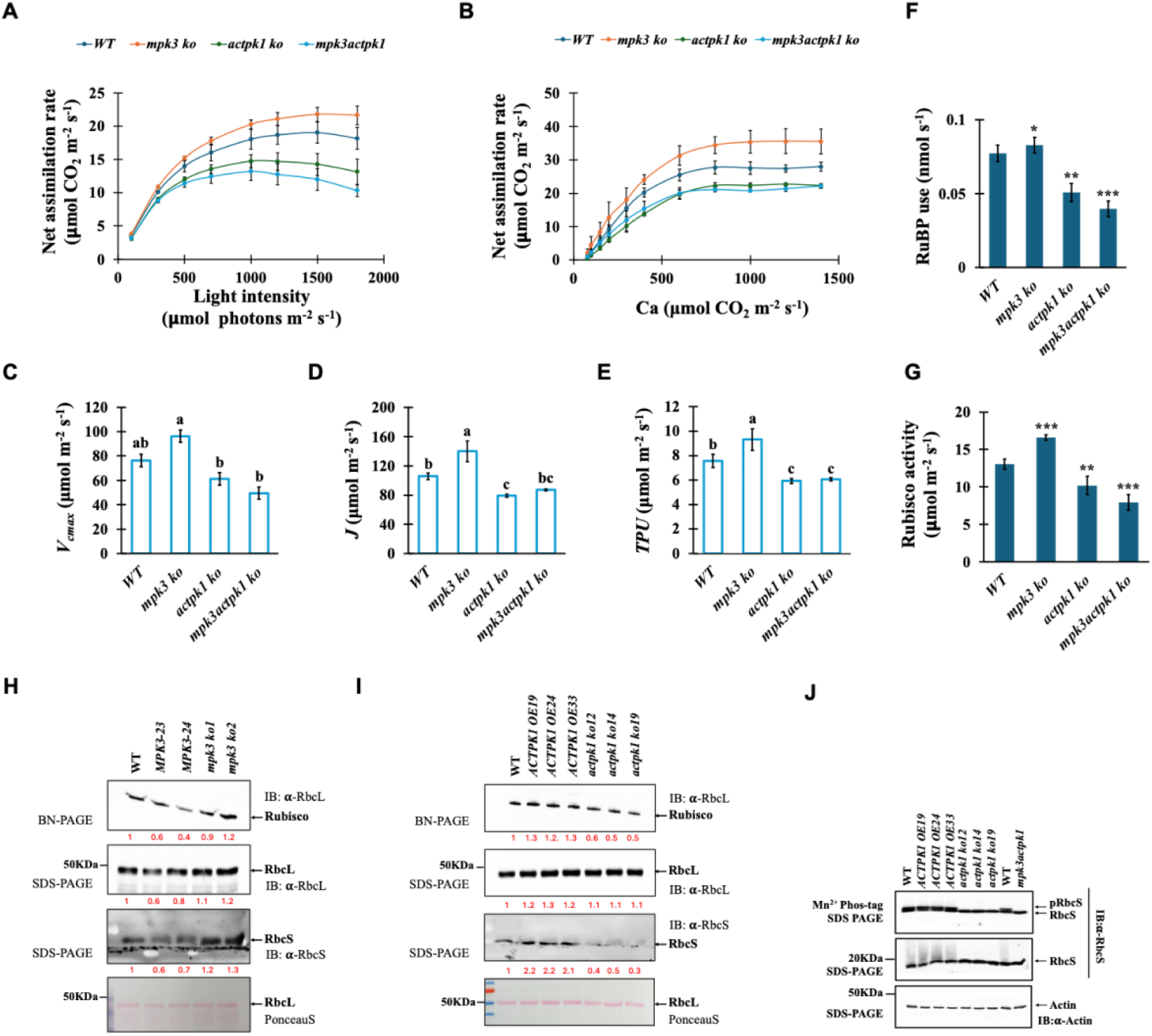
ACTPK1 acts downstream of MPK3 to regulates photosynthesis in rice. **A)** Comparison of net photosynthesis per unit area of WT, *mpk3 ko*, *actpk1 ko* and *mpk3actpk1 ko* plants in response to varying CO_2_ concentration at 1500 μmol m^−2^ s^−1^ light intensity (*A/Ci* curve). Each value represents mean ± SD, where n = at least three data points from different plants. **B)** Comparison of net photosynthesis per unit area of WT, *mpk3 ko*, *actpk1 ko* and *mpk3actpk1 ko* plants in response to increasing PPFD at 400 μmol mol^−1^ CO_2_ concentration (*A/Q* curve). Each value represents mean ± SD, where n = at least three data points from different plants. **C)** Bar graph showing maximum rate of carboxylation (*Vcmax*) in WT, *mpk3 ko*, *actpk1 ko* and *mpk3actpk1 ko* plants. **D)** Bar graph showing electron transport rate (*J*) in of WT, *mpk3 ko*, *actpk1 ko* and *mpk3actpk1 ko* plants. **E)** Bar graph showing triose phosphate utilization (*TPU*) in of WT, *mpk3 ko*, *actpk1 ko* and *mpk3actpk1 ko* plants. Data represented in D), E) and F) are deduced by fitting *A/Ci* curve using FvcB model. All the measurements were taken from flag leaf at heading stage using LiCOR6800, between 8:00 am to 11:00 am. Different letters indicate statistical significance according to one-way ANOVA followed by post-hoc Tukey HSD calculation at *P < 0.05*. **F)** Bar graph showing RuBP use in in WT, *ACTPK1 OE* (*ACTPK1 OE19, ACTPK1 OE24 & ACTPK1 OE33*) and *actpk1 ko* (*actpk1 ko12, actpk1 ko14 & actpk1 ko19*) and *mpk3actpk1 ko* lines. **G)** Bar graph showing Rubisco activity in WT, *ACTPK1 OE* (*ACTPK1 OE19, ACTPK1 OE24 & ACTPK1 OE33*) and *actpk1 ko* (*actpk1 ko12, actpk1 ko14 & actpk1 ko19*) and *mpk3actpk1 ko* lines. Values represent the mean ± SD (*n* = 3 biological replicates). Asterix indicate statistical significance according to student ttest (*P*<0.05). **H)** Immunoblots showing Rubisco (upper panel), RbcL (middle panel) and RbcS (lower panel) content in WT, *MPK3 OE* (*MPK3-23 & MPK3-24*) and mpk3 ko (*mpk3 ko1 & mpk3 ko2*) plants. Blots were probed with α-RbcL antibody for Rubisco and RbcL and with α-RbcS antibody for RbcS content. **I)** Immunoblots showing Rubisco (upper panel), RbcL (middle panel) and RbcS (lower panel) content in in WT, *ACTPK1 OE* (*ACTPK1 OE19, ACTPK1 OE24 & ACTPK1 OE33*) and *actpk1 ko* (*actpk1 ko12, actpk1 ko14 & actpk1 ko19*) plants. Blots were probed with α-RbcL antibody for Rubisco and RbcL and with α-RbcS antibody for RbcS content. Band intensities (written in red color) in (H) and (I) were quantified using ImageJ software. **J)** Phos-tag gel western blots showing phosphorylated and non-phosphorylated RbcS content in WT, *ACTPK1 OE* (*ACTPK1 OE19, ACTPK1 OE24 & ACTPK1 OE33*) and *actpk1 ko* (*actpk1 ko12, actpk1 ko14 & actpk1 ko19*) and *mpk3actpk1 ko* (double mutant) lines. Blots were probed with α-RbcS antibody. Actin was used as endogenous control.

Moreover, the biochemical insights support the above observations. *Vcmax* was highest in *mpk3 ko* lines but drops to *actpk1 ko* levels in and *mpk3actpk1 ko* plants*. J* and *TPU* followed the same trend, enhanced in *mpk3 ko*, and reduced in *actpk1 ko* and *mpk3actpk1 ko* plants compared to WT. (Fig. 6C, D & E). Rubisco activity measurements further confirmed this trend, with higher activity in *mpk3 ko* and strongly diminished activity in *actpk1 ko* and *mpk3actpk1 ko* (Fig. 6F, G).

Confirming the observation of the altered carboxylation capacity, BN-PAGE and immunoblot analyses revealed that Rubisco holoenzyme abundance, as well as RbcS protein levels, were increased in *mpk3 ko* plants and reduced in *MPK3 OE* lines compared with WT (Fig. 6H). Conversely, *actpk1 ko* plants showed reduced Rubisco holoenzyme abundance together with decreased RbcS levels, whereas *ACTPK1 OE* plants accumulated higher amounts of Rubisco and RbcS relative to WT (Fig. 6I). These changes in Rubisco accumulation mirror the corresponding alterations in photosynthetic capacity across the MPK3 and ACTPK1 genetic backgrounds.

The finding that *mpk3actpk1* double mutants phenocopy the *actpk1 ko* mutants, rather than *mpk3 ko* alone, indicates that ACTPK1 acts genetically downstream of MPK3. Moreover, the loss of the enhanced photosynthetic performance observed in *mpk3 ko* plants upon ACTPK1 disruption highlights a linear regulatory hierarchical, in which MPK3 negatively regulates photosynthesis by suppressing ACTPK1 activity.

To directly link the genetic hierarchy to RbcS regulation, we examined RbcS phosphorylation across these genotypes using Phos-tag SDS–PAGE. Phos-tag analysis revealed a distinct mobility shift of RbcS in WT and *ACTPK1 OE* seedlings, indicative of phosphorylation. In contrast, no shift was detected in *actpk1 ko* or *mpk3actpk1 ko* seedlings, demonstrating that enhanced RbcS phosphorylation in *mpk3 ko* plants is due to the presence of ACTPK1, thus placing ACTPK1 downstream in this cascade (Fig. 6J). This hierarchy aligns with our biochemical evidence that MPK3 phosphorylates ACTPK1 thereby inhibiting its kinase activity and preventing ACTPK1-mediated phosphorylation of RbcS.

### ACTPK1 positively regulates plant biomass and yield

Photosynthetic efficiency directly influences plant productivity, where enhanced carbon assimilation supports plant growth and biomass. We next examined the effect of ACTPK1-mediated photosynthesis regulation on plant growth and development. *ACTPK1 OE* and WT seedlings exhibited enhanced shoot and root length compared to *actpk1 ko* which displayed retarded growth, indicating robust vegetative growth fueled by increased carbon fixation (Supplemental Fig. S9A-C). At maturity, plant height, tiller number per plant, and plant biomass were significantly reduced in *actpk1 ko* lines relative to *ACTPK1 OE* and WT (Fig. 7A-D). Moreover, striking differences were observed in flag leaf angle, *ACTPK1 OE* plants displayed highly erect flag leaves with narrow angles (8°–19°), WT plants had moderately open leaves (14°–29°), whereas *actpk1 ko* lines exhibited leaves with widest angles (45°–75°) (Supplemental Fig. S9D-E). As leaf angle strongly influences light interception, these differences likely have direct consequences for photosynthetic efficiency. Furthermore, flowering time was markedly delayed in *actpk1 ko* by 10–15 days compared to *ACTPK1 OE* and WT. While WT and *ACTPK1 OE* lines-initiated flowering at 80–85 days, *actpk1 ko* plants flowered at approximately 100–105 days after sowing, indicating a potential regulatory role of ACTPK1 in floral transition (Supplemental Fig. S9F). These data suggest that absence of *ACTPK1* disrupts developmental timing and architecture, likely due to impaired photosynthesis and resource allocation.

**Fig. 7.**
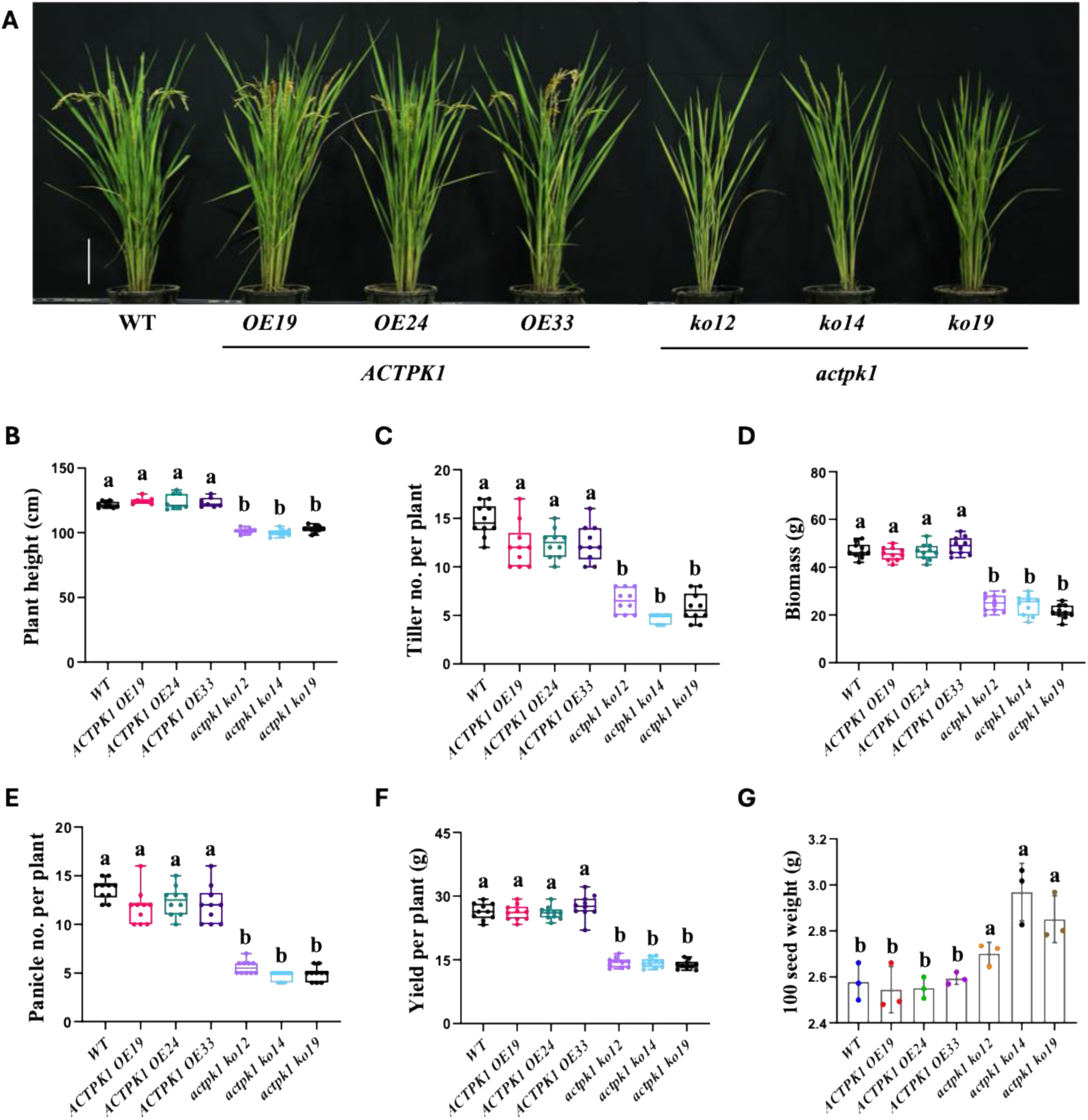
ACTPK1 positively regulate plant biomass and yield. **A)** Representative image of WT, *ACTPK1 OE* (*ACTPK1 OE19, ACTPK1 OE24 & ACTPK1 OE33*) and *actpk1 ko* (*actpk1 ko12, actpk1 ko14 & actpk1 ko19*) plants. **B)-G)** Bar graph showing plant height **(B)**, tiller number per plant **(C)**, Biomass **(D)**, panicle number per plant **(E)**, yield per plant **(F)**, and 100 seed weight **(G)**, in WT, *ACTPK1 OE* (*ACTPK1 OE19, ACTPK1 OE24 & ACTPK1 OE33*) and *actpk1 ko* (*actpk1 ko12, actpk1 ko14 & actpk1 ko19*) plants. Values represent the mean ± SD (*n* = 10). Different letters indicate statistical significance according to one-way ANOVA followed by post-hoc Tukey HSD calculation at *P < 0.05*.

Interestingly, *ACTPK1* also affects seed architecture and yield. *actpk1 ko* plants exhibited significantly lower number of panicles per plant compared to *ACTPK1 OE* and WT (Fig. 7E). Lower panicle number resulted in reduced grain yield per plant in *actpk1 ko* plants (Fig. 7F). Interestingly, seeds from *actpk1 ko* plants were not only longer but also wider than those from WT and *ACTPK1* OE lines, resulting in a significantly higher 100-seed weight (Fig. 7G, Supplemental Fig. S9G-I). However, these advantages in seed productivity were offset by a substantial reduction in tiller number and panicle number, traits that are key contributors to overall yield. Thus, while *ACTPK1* knockout promotes the formation of larger and heavier seeds, it concurrently suppresses the production of structures such as tillers and panicles that bears reproductive organs, highlighting its role in balancing the trade-off between seed size and seed number.

## Discussion

Chloroplast protein import must be tightly coordinated with cytosolic signalling to sustain photosynthetic performance. Although phosphorylation of chloroplast transit peptides has been proposed to influence precursor targeting for more than two decades, the physiological relevance and regulatory logic of this modification have remained unresolved. Here, we identify a MAPK-regulated signalling module in rice in which MPK3 modulates the activity of the cytosolic Raf-like kinase, ACTPK1 to dynamically regulate the phosphorylation status of the RbcS precursor. Our results demonstrate that neither constitutive phosphorylation nor permanent loss of phosphorylation is sufficient for efficient RbcS import, indicating that reversible phosphorylation rather than a fixed phosphorylation state underpins productive chloroplast translocation. This dynamic regulatory model reconciles earlier biochemical observations with *in vivo* import behaviour and provides a mechanistic framework through which cytosolic signalling pathways can fine-tune chloroplast protein targeting and photosynthetic capacity.

The central mechanistic finding of this study is that MPK3 negatively regulates ACTPK1, which directly phosphorylates the RbcS precursor at Thr12 within its transit peptide. Genetic and biochemical analyses demonstrate that MPK3 suppresses ACTPK1 activity, thereby modulating RbcS phosphorylation, Rubisco accumulation and photosynthetic capacity.

Phosphorylation has long been implicated in chloroplast protein import, particularly through modification of transit peptides that modulate precursor handling and targeting. In Arabidopsis, the STY kinases STY8, STY17 and STY46 phosphorylate RbcS transit peptides and influence chloroplast protein import (Lamberti et al., 2011). Our identification of ACTPK1 as the functional rice homolog of this kinase family places RbcS transit-peptide phosphorylation within a genetically defined MPK3-regulated signaling pathway.

MPK3, localized in the cytoplasm and nucleus, indirectly regulates photosynthesis by modulating RbcS phosphorylation dynamics through ACTPK1. Protein phosphorylation is a well-established regulator of photosynthesis, coordinating light harvesting, energy distribution and photoprotection (Tikkanen and Aro, 2012; Rochaix, 2014), and contributes to chloroplast protein import by modifying transit peptides and translocon components (Waegemann and Soll, 1996; May and Soll, 2000; Jarvis and Robinson, 2004). Rubisco subunits themselves are known phosphoproteins, and these modifications can influence enzyme stability and function (Houtz et al., 1989; Grabsztunowicz et al., 2017). In accordance with this regulatory framework, we show that RbcS phosphorylation is enhanced in *mpk3 ko* plants and is associated with increased Rubisco accumulation and elevated carboxylation efficiency. Mechanistically, ACTPK1 directly phosphorylates the RbcS precursor, thereby providing the molecular basis through which MPK3 signaling regulates chloroplast protein targeting.

MPK3 directly phosphorylates ACTPK1, providing a mechanistic basis for its negative regulation and establishing a non-canonical MAPK module linking cytosolic signalling to chloroplast protein targeting. Protein–protein interaction assays, including Y2H, BiFC and Co-IP, demonstrate direct association between MPK3 and ACTPK1, while *in vitro* kinase assays establish unidirectional phosphorylation of ACTPK1 by MPK3. Functional analyses further show that ACTPK1 kinase activity is enhanced in *mpk3 ko* plants, demonstrating that MPK3-mediated phosphorylation negatively regulates ACTPK1 activity. This kinase-on-kinase regulation explains how MPK3 modulates RbcS phosphorylation, Rubisco accumulation and photosynthetic performance.

ACTPK1 loss also causes pleiotropic developmental and yield-related phenotypes, including delayed flowering, reduced tillering and altered leaf architecture, which correlate with reduced CO₂ assimilation and carboxylation efficiency. These findings indicate that ACTPK1 activity is required to sustain photosynthetic capacity and normal plant productivity. Given the established role of ACTPK1 in ammonium transporter regulation (Beier et al., 2018; Shilpha et al., 2023), these phenotypes likely reflect combined effects of impaired photosynthesis and altered metabolic homeostasis rather than a direct developmental signaling function of the MPK3–ACTPK1–RbcS module.

At the molecular level, ACTPK1 phosphorylates RbcS at Thr12 within its transit peptide, as demonstrated by *in vitro* kinase assays and site-directed mutagenesis. This modification contributes to maintaining RbcS in an import-competent cytosolic state. On the other hand our phosphomutant and inhibitor analyses indicate that reversible phosphorylation, finely coordinating kinase and phosphatase activities, is necessary to support efficient chloroplast translocation. In accordance with this model, the phosphomimetic RbcS^T12D^ variant and phospho-dead RbcS^T12A^ variant shows reduced chloroplast targeting, in agreement with previous studies demonstrating the importance of transit peptide phosphorylation dynamics for precursor handling and import (Waegemann and Soll, 1996; May and Soll, 2000; Lee et al., 2009; Jarvis & López-Juez, 2013). These observations indicate that phosphorylation is not simply required or inhibitory for import, but rather that regulated phosphorylation dynamics are necessary to maintain precursor competence for chloroplast targeting. Transit peptide phosphorylation has been proposed to regulate precursor interaction with cytosolic chaperones such as 14-3-3 proteins and Hsp70, which maintain precursors in an import-competent state, suggesting that ACTPK1-mediated phosphorylation may regulate RbcS through similar mechanisms.

The absence of detectable RbcS–GFP signal in *actpk1* mutant cells, together with its rapid destabilization in degradation assays, suggests that ACTPK1 activity contributes to maintaining RbcS precursor accumulation, likely by promoting precursor stability and import competence. Because MG132 treatment did not restore RbcS–GFP accumulation, the precise degradation pathway remains unclear, suggesting that multiple quality-control mechanisms may contribute to precursor turnover in the absence of ACTPK1. These findings support a model in which ACTPK1-dependent phosphorylation contributes to maintaining precursor accumulation and import competence, which in turn facilitates efficient Rubisco assembly and photosynthetic function. In line with this interpretation, increased ACTPK1 activity in *mpk3* knockout plants correlates with enhanced Rubisco accumulation and photosynthetic capacity, whereas loss of ACTPK1 leads to reduced Rubisco levels and impaired photosynthetic performance. However, the molecular mechanisms responsible for RbcS destabilization in the absence of ACTPK1 remain to be elucidated, and future studies will be required to identify the quality-control pathways that regulate precursor turnover. In addition, reduced RbcS transcript levels in actpk1 mutants indicate that ACTPK1 may influence RbcS abundance through both post-transcriptional and transcriptional mechanisms.

Genetic epistasis analysis establishes a linear signalling hierarchy in which ACTPK1 functions downstream of MPK3. The *mpk3actpk1* double mutant phenocopies *actpk1* knockout plants in terms of RbcS phosphorylation, Rubisco activity and photosynthetic capacity, demonstrating that ACTPK1 is required for the photosynthetic effects of MPK3. Based on our findings under non-stress growth conditions, we propose that the MPK3–ACTPK1 module may provide a regulatory interface through which signalling pathways could modulate RbcS handling and Rubisco biogenesis. Although the upstream signals that engage MPK3 in this context remain unknown, it is plausible that modulation of MPK3 activity could influence ACTPK1-dependent RbcS phosphorylation and thereby tune photosynthetic capacity.

Several mechanistic aspects of the MPK3–ACTPK1–RbcS pathway remain unresolved. Although our data demonstrate that dynamic phosphorylation of the RbcS transit peptide is required for efficient chloroplast targeting, the phosphatase responsible for RbcS dephosphorylation has not yet been identified. In addition, while loss of ACTPK1 leads to rapid loss of detectable RbcS and accelerated degradation in cell-free assays, the molecular components of the cytosolic quality-control pathway responsible for RbcS turnover in the absence of ACTPK1 remain unknown. Furthermore, although ACTPK1 directly phosphorylates RbcS at Thr12 in vitro and genetic analyses support its role in vivo, the precise molecular mechanism by which phosphorylation modulates precursor recognition, chaperone engagement and TOC/TIC translocation remains to be defined. Finally, whether ACTPK1 regulates additional chloroplast-destined precursor proteins beyond RbcS is currently unknown and will require systematic proteomic and genetic analyses.

In summary, our study reveals a signaling mechanism through which cytosolic MAPK activity influences chloroplast protein import and photosynthetic performance. We identify ACTPK1 as a downstream target of MPK3 and show that this kinase regulates the stability and chloroplast targeting competence of the Rubisco small subunit precursor through phosphorylation of its transit peptide. Our results further indicate that efficient handling of the RbcS precursor requires a dynamically regulated phosphorylation state rather than a constitutive modification, highlighting transit peptide phosphorylation as a reversible regulatory element in chloroplast protein trafficking. A simplified model of the findings has been represented in Figure 8. Together, these findings uncover a signaling axis that links cytosolic MAPK pathways with the control of Rubisco biogenesis and photosynthetic capacity, providing a framework for understanding how cellular signaling networks coordinate chloroplast function with broader physiological demands in plants.

**Fig. 8.**
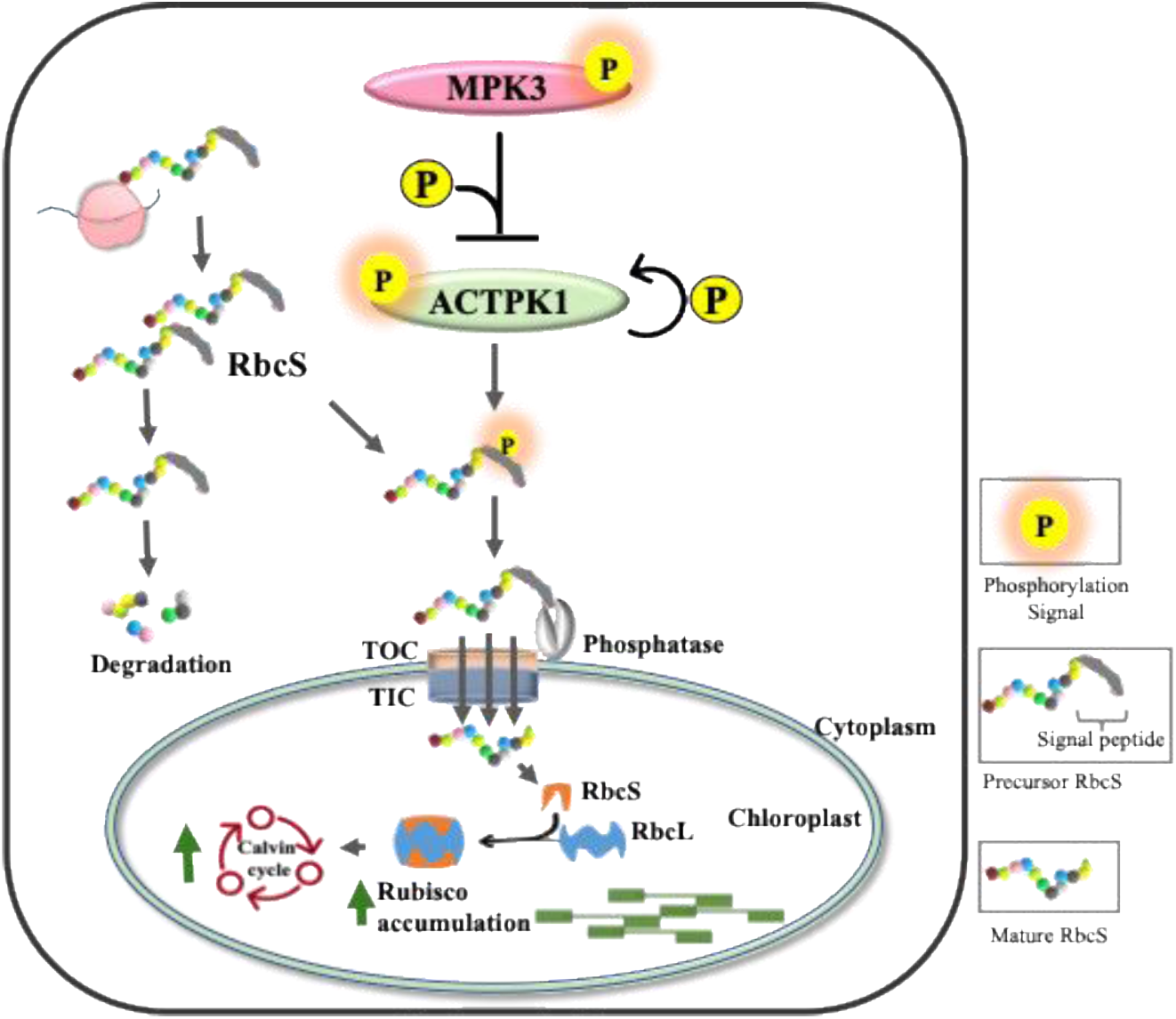
A MAPK signaling module linking cytoplasmic signaling to Rubisco biogenesis. In this model, MPK3 phosphorylates and inhibits the Raf-like kinase ACTPK1. ACTPK1 directly phosphorylates the Rubisco small subunit precursor (RbcS) at Thr12 within its transit peptide, stabilizing the precursor and increasing its import competence into chloroplasts. Through this mechanism, the MPK3–ACTPK1–RbcS signaling cascade connects cytoplasmic kinase signaling with chloroplast protein biogenesis to fine-tune Rubisco accumulation and photosynthetic carbon fixation.

## Materials and Methods

### Transgene construct

For generation of overexpression lines, full-length coding sequence (CDS) of *ACTPK1* was amplified with 3xFLAG tag at 5’ end and 4xMYC tag at 3’ end was sandwiched between 35S promoter and Nos terminator and cloned into binary vector pCAMBIA1300-CaMV 35S. The pCAMBIA1300-35S-*ACTPK1* plasmid was introduced into rice *japonica* cultivar Taipea309 (Tp309) via *Agrobacterium*-mediated transformation using *Agrobacterium* strain EHA105.

*ACTPK1* knockout mutants were generated using CRISPR-Cas9 mediated gene editing in japonica cultivar Tp309 background according to Rengasamy et al., 2024. A single guide RNA (sgRNA) targeting the CDS region of *ACTPK1* gene (sgRNA: GGGGACGGCGCGCGGACAAGAGC, PAM sequence: GGG) was designed and synthesized as an oligonucleotide. The synthesized sgRNA was then cloned in pRGB32 vector and introduced into 10-day-old rice Calli through *Agrobacterium*-mediated transformation. The *MPK3-ACTPK1* double knockout was generated by mutating *ACTPK1* in *mpk3 ko* background using CRISPR-Cas9 genome editing tool using the same sgRNA (sgRNA: GGGGACGGCGCGCGGACAAGAGC).

### Plant growth conditions

Rice seeds used for photosynthesis experiments were homozygous seeds of atleast T2 generation. Along with *ACTPK1 OE*, *actpk1 ko* and *mpk3-actpk1* double knockout, dexamethasone (DEX) inducible MPK3 OE lines (*MPK3-23*, *MPK3-24*) and *mpk3 ko* (*mpk3 ko1*, *mpk3 ko2*) plants (both generated in *Oryza sativa* Tp309 cultivar) were used (Singh et al 2019, Banerjee et al., 2025). Seeds were germinated in petri plates having moist germination paper for 7 days at 28 °C, 16h light and 8h dark conditions. Seven days old seedlings were transferred to one-fourth M.S. (Himedia) hydroponics media containing both micro and macronutrients. The pH of the media solution was adjusted to 5.5, and media was renewed every 3 days. Post 20 days, plants were transferred to pots and kept in a greenhouse at 28/22 °C day and night air temperature, 70% relative humidity, a photoperiodic cycle of 16h light/8h dark photoperiod, and ∼500 µmol m^−2^ s^−1^ light intensity. The plants were maintained in pots in greenhouse until maturity, and photosynthetic parameters were recorded from the middle widest area of flag leaf at heading stage. For *MPK3 OE* lines, 1 um DEX and DMSO (mock) were sprayed one day prior to taking photosynthetic readings (Singh et al., 2019).

### Photosynthesis rate and related physiological trait measurements

Photosynthetic rate and its related physiological parameters were measured using Li-COR 6800 portable photosynthesis system on the flag leaf of rice plant at heading stage. The area of Li-COR 6800 portable system leaf chamber was set at 2 cm^2^ with a LED light source (LI-6400-02B; LI-COR, Inc). To acquire the same leaf area, two young leaves were carefully placed in leaf chamber without any overlap for plants having leaf width less than 2 cm^2^ (Busch et al, 2018). For all the measurements, the conditions of leaf chamber were set as follows: 1500 µmol m^−2^ s^−1^ light intensity (PAR), 400 μmol mol^−1^ CO_2_ concentration, 300 µmol s^−1^ constant airflow, 70-80% relative humidity. Ten flag leaves from different plants were used to evaluate the net photosynthesis rate (*A*), intercellular CO_2_ concentration (*Ci*) and stomatal conductance to water (*gs*).

For A/Ci curve, the leaves were first acclimatised to ambient CO_2_ concentration of 400 μmol mol^−1^ CO_2_ and a constant PAR of 1500 μmol m^−2^ s^−1^ for 10 min and then the CO_2_ concentration was first gradually decreased from 400 to 50 (400, 300, 200, 150, 100, 50) μmol mol^−1^ CO_2_ and then leaf was stabilized at 400 μmol mol^−1^ CO_2_ and again CO_2_ concentration was increased from 400 to 1400 (400, 600, 800, 1000, 1200 and 400) μmol mol^−1^ CO_2_. The minimum and maximum wait time between the subsequent readings was 60 and 120 sec respectively. Readings were taken at constant PAR of 1500 μmol m^−2^ s^−1^, flow rate of 500 µmol s^−1^. The maximum rate of Rubisco mediated carboxylation (*Vcmax*), electron transport rate (*J*) and triose phosphate utilisation (*TPU*) were calculated by fitting A/Ci curve using Farquhar-von Caemmerer-Berry (FvcB model) (Sharkey, 2015).

For light curve, the leaf was first acclimatised to constant saturating photosynthetic active radiation (PAR) of 1500 μmol m^−2^ s^−1^ for about 10 min and then the PAR was decreased from 1500 to 100 (1500,1200,1000, 700, 500, 300 and 100) μmol m^−2^ s^−1^ at constant CO_2_ concentration of 400 μmol mol^−1^ CO_2_ and flow rate of 500 µmol s^−1^. The minimum and maximum wait time between the subsequent readings was 120 and 180 sec respectively.

All measurements were taken between 9.00 am to 11.00 am.

### Gene expression analysis

Total RNA was isolated from leaf samples using RNeasy Plant mini kit (Qiagen) following manufacturer’s protocol. DNase treatment was given to 2 μg RNA using RNA free DNase I (Qiagen) and subjected to first strand cDNA synthesis using RevertAid First Strand cDNA synthesis kit (Thermo Scientific) per the manufacturer’s protocol. Quantitative RT-PCR of ACTPK1 and RbcS was carried out using Power SYBR Green PCR master mix (Applied Biosystems). The conditions of thermocycler were set as follows: 94 °C for 10 min, 40 cycles of 94 °C for 20 sec, 60 °C for 1 min. The housekeeping gene UBQ5 and Actin7 were used as an internal control for normalization. Expression levels of examined genes were quantified using relative quantitation method (ΔΔCT). The statistical significance was evaluated using t-test. Data shown is the mean value of at least three biological repeats with SD. The primers used for doing qRT-PCR are listed in supplemental Table 1.

### Site-directed mutagenesis

Site-directed mutagenesis was performed using overlap extension PCR. Complementary primers containing the desired mutation were designed. The target gene was amplified in two fragments using gene-specific forward and mutation-specific reverse primers, and mutation-specific forward and gene-specific reverse primers. Both amplicons were gel-purified and used in overlap extension PCR with the following conditions: 98°C for 30 s, 55–58°C for 1 min, 72°C for 30 s/kb for 10 cycles. Gene-specific primers were then added for 35 additional cycles. The final PCR product was gel-purified, verified by DNA sequencing, and cloned into the target vector.

### Bacterial protein induction, purification and *in vitro* phosphorylation assay

The full length CDS of *MPK3*, *MPK6*, *ACTPK1*, and *RbcS* were cloned in-frame with the sequence of GST-tag in pGEX4-T-2 vector. Recombinant protein was induced in *Escherichia coli BL21 Rosetta*^®^ by adding 1mM IPTG at 22 °C. Induced recombinant proteins with GST-tag were purified by affinity chromatography using GST-sepharose beads. Protein concentration was determined using Bradford reagent (Sigma) and purity was analysed using SDS-PAGE. *In vitro* phosphorylation assay was performed according to the method described by Raghuram *et al*, 2015. Briefly, purified recombinant proteins of kinase and substrate (ratio 1:10) were incubated in kinase reaction buffer (25 mM Tris/Cl (pH 7.5), 10 mM MgCl_2_, 5 mM MnCl_2_, 1 mM DTT, 1 mM β-glycerol phosphate, 1 mM Na_3_VO_4_, 25 mM ATP and 1 mCi [γ-32P] ATP) for 30 min at 30 °C. The reaction was stopped by adding 6X SDS sample buffer followed by heating at 95 °C for 5 min. The reaction samples were subjected to 12% SDS-PAGE and exposed to phosphor screen followed by visualization using phosphor imager (Typhoon, Phosphor Imaging System, GE Health Care, Life Sciences).

### Plant protein extraction and Immunoblot Analysis

For SDS-PAGE, Proteins were extracted from rice seedlings using protein extraction buffer (50 mM Tris HCl (pH 7.5), 5 mM EDTA, 5 mM EGTA, 150 mM NaCl, 1 mM DTT, 10 mM

Na_3_VO_4_, 10 mM NaF, 1mM PMSF, 2 mg/ml PVPP, 10% glycerol (v/v), 0.1% Nonidet P-40 (v/v) and 1× protease inhibitor cocktail (Sigma-Aldrich/Merck)). Extracted proteins were quantified with Bradford reagent and ∼40 μg was separated by 10/12% SDS-PAGE. Specifically, 1 and 5 μg protein was resolved for RbcL and RbcS content quantification respectively. Separated proteins were transferred to PVDF membrane for immunodetection. The membranes were first blocked with 5% skimmed milk in TBST (10 mM Tris (pH 7.5), 500 mM NaCl, 0.05% Tween 20) for 2 h, then probed with anti RbcS (Agrisera, dilution 1:5000), anti Actin (Sigma-Aldrich, dilution 1:5,000) antibody in TBST for 2 h at room temperature. After two TBST washes (2 min each), membranes were incubated with anti-rabbit secondary antibody (Thermofisher Scientific A11034) for 1.5 h. The membrane was again washed twice with TBST (10 min each) and MQ water and visualized using Immobilon Western chemiluminescent HRP substrate (BIORAD) as per manufacturer’s instructions. Chemiluminescence was detected with ChemiDoc system iBright 1500 (Amersham Biosciences).

For quantification of Rubisco, Proteins were extracted using non-denaturing extraction buffer (50 mM Tris HCl (pH 7.5), 50 mM NaCl, 1 mM DTT, 2 mg/ml PVPP, 10% glycerol (v/v), 0.1% Nonidet P-40 (v/v) and 1× protease inhibitor cocktail (Sigma-Aldrich/Merck). 1 μg protein was resolved on 5-15% gradient BN-PAGE, transferred to PVDF membrane and probed with anti-RbcL antibody.

### *In vivo* phosphorylation assay

*In vivo* phosphorylation of RbcS was examined by mobility shift assay employing phos-tag reagent. Proteins were extracted from WT, *mpk3 ko, ACTPK1 OE, actpk1 ko, mpk3actpk1 ko* seedlings using protein extraction buffer (50 mM Tris HCl (pH 7.5), 150 mM NaCl, 1 mM DTT, 10 mM NaF, 1mM PMSF, 10% glycerol (v/v) and 0.1% Nonidet P-40 (v/v)). Extracted protein was separated by 12% (w/v) SDS-PAGE gel containing 100 μM phos-tag and 100 mM MnCl_2_. After electrophoresis, the gel was washed thrice, twice with transfer buffer containing 10 mM EDTA and once without EDTA for 10 min. Post washing, separated proteins were transferred to nitrocellulose membrane using Trans-Blot SD cell (BIO-RAD) and membrane was probed with α-RbcS (Agrisera) antibody. The membrane was again washed twice with TBST (10 min each) and MQ water and visualized using Immobilon Western chemiluminescent HRP substrate (BIORAD) as per manufacturer’s instructions. Chemiluminescence was detected with ChemiDoc system iBright 1500 (Amersham Biosciences).

### Immuno-kinase (IP-kinase) assay

Total protein was extracted from the 14 day old seedlings of WT, *mpk3 ko1* and *mpk3 ko2* using the previously described protein extraction. Extracted proteins were quantified using Bradford reagent, and 1 mg crude protein was incubated with 5 μg anti ACTPK1 antibody (genescript) on a rocker at 4°C for 3 h. Subsequently, 50 μL Protein A Sepharose beads (50% slurry, Pierce) were added and incubated on a rocker at 4°C for 1.5 h. Post incubation, sepharose beads were washed with PBS-T (PBS with 0.05% Tween 20) thrice, resuspended in kinase buffer containing 1 mCi [γ-³²P] ATP and 0.5 μg MBP, and incubated at 30°C for 30 min. Reactions were stopped with 3× SDS dye, boiled at 95°C for 5 min, and resolved by 12% SDS-PAGE. Gels were exposed to a phosphor screen and visualized using a Typhoon phosphor imager (GE Healthcare).

### Yeast two-hybrid assay

The interaction between MPK3 and ACTPK1, and ACTPK1 and RbcS was examined using the Matchmaker Gold Yeast Two-hybrid System (Takara Bio). Full-length CDS of MPK3, MPK6, ACTPK1 and RbcS were cloned in pGBKT7 and pGADT7 vector (Clontech). Yeast transformation with was performed using Fast Yeast Transformation kit (Gbiosciences; GZ-1) (according to the manufacturer’s protocol). The positive transformants were selected on double dropout medium (Synthetic Defined (SD) /-Leu/-Trp)). The interaction assay was performed by plating the selected transformants on quadruple dropout medium ((SD)/-Ade/-His/-Leu/-Trp) followed by incubation at 30 °C for 36–48h. The fully grown colonies after the incubation were marked as positive interactions.

### Subcellular localisation and Bimolecular Fluorescence Complementation assay (BiFC)

For subcellular localisation, full-length CDS of MPK3 and RbcS was cloned in pGWB5 vector. The resulting construct (MPK3-pGWB5) was transformed into *Agrobacterium* strain EHA105 and infiltrated in young and healthy leaves of *Nicotiana benthamiana*. The plasmid of RbcS-pGWB5, RbcS^T12D^-pGWB5 and RbcS^T12A^-pGWB5 was introduced in rice protoplast using PEG-mediated transformation (citation). Localisation was analysed post 48h for *Nicotiana benthamiana* and post 16h for rice protoplast using a Leica TCS SP8 AOBS Laser Scanning Microscope (Leica Microsystems) using GFP filters with fluorescence excitation at 514 nm and emission at 527 nm.

For BiFC, full length CDS of MPK3, ACTPK1 and RbcS was cloned into pSITE-nEYFP-C1 (Cd3-1648) and pSITE-cEYFP-N1 (Cd3-1651) vectors. The resulting constructs were transformed into *Agrobacterium* strain EHA105 and infiltrated in young and healthy leaves of *Nicotiana benthamiana*. Interaction was analysed post 48h using a Leica TCS SP8 AOBS Laser Scanning Microscope (Leica Microsystems) using YFP filters with fluorescence excitation at 514 nm and emission at 527 nm.

### Co-immunoprecipitation (Co-IP) assay

Flag-tagged ACTPK1 (pCAMBIA1300-35S-Flag-*ACTPK1*) and GFP-tagged MPK3 (MPK3-pGWB5) were co-expressed in *Nicotiana benthamiana* leaves via *Agrobacterium*-mediated infiltration. Proteins were extracted 48 h post-infiltration using protein extraction buffer. Proteins (1 mg) were incubated with 5 μg anti-Flag antibody (Sigma-Aldrich) at 4°C for 3 h with rocking, followed by 50 μL Protein A Sepharose beads (50% slurry, Pierce) for 1.5 h. Beads were washed thrice with PBS-T (PBS, 0.05% Tween 20), eluted with 2× SDS sample buffer, boiled at 95°C for 5 min, and resolved on 10% SDS-PAGE. Proteins were transferred to PVDF, blocked with 5% skimmed milk in TBST (10 mM Tris, pH 7.5; 500 mM NaCl; 0.05% Tween 20) for 2 h, and probed with anti-GFP antibody (Abcam) for 2 h at room temperature. After two 10-min TBST washes, membranes were incubated with HRP-conjugated secondary antibody for 1.5 h, washed twice with TBST and MQ water, and visualized using Immobilon Western chemiluminescent HRP substrate (BIORAD) as per manufacturer’s instructions. Chemiluminescence was detected with ChemiDoc system iBright 1500 (Amersham Biosciences).

### Cell-free degradation assay

Total protein was extracted from 14-day-old rice seedlings (WT, *mpk3 ko*, *actpk1 ko*) by grinding 1 g tissue in liquid nitrogen and homogenizing in 1 mL ice-cold protein extraction buffer. Extracted proteins were quantified using Bradford reagent. Crude extract (100 µg) was incubated with 1 µg recombinant GST-tagged RbcS in 100 µL reaction buffer (50 mM Tris-HCl, (pH 7.5), 5 mM MgCl₂, 5 mM ATP, and 1 mM DTT) at room temperature. Aliquots (20 µL) were collected at 0, 5, 10, 30, 60, and 120 min, stopped with 2× SDS sample buffer, boiled at 95°C for 5 min, and resolved on 12% SDS-PAGE. Proteins were transferred to nitrocellulose membrane and probed with anti-GST antibody (Thermofischer), and visualized using Immobilon Western chemiluminescent HRP substrate (BIORAD) as per manufacturer’s instructions. Chemiluminescence was detected with ChemiDoc system iBright 1500 (Amersham Biosciences). Band intensity was quantified with ImageJ.

### Rubisco activity measurements

Rubisco activity assay was performed according to Sharwood et al., (2016). Briefly, Leaf tissues were excised (2 cm^2^) from the middle widest part of the same leaf from which photosynthesis parameters were recorded, pulverized and homogenized in 1 mL ice-cold extraction buffer (5 mM DTT, 10 mg ml^−1^ PVPP, 2 mg ml^−1^ BSA, 2 mg ml^−1^ PEG, 2% (v/v) Tween-80, 12 mM 6-aminocaproic acid, 2.4 mM benzamidine in HEPES stock) at 4 °C. The homogenate was centrifuged at 15,000 × g at 4°C for 1 min, and supernatant was incubated with freshly prepared BMEC buffer (100 mM Bicine–NaOH, pH 8.2, 20 mM MgCl_2_, 1 mM Na_2_-EDTA, and 10 mM NaHCO_3_) in a 9:1 ratio (e.g. 900 μl leaf extract to 100 μl BMEC) at 25 °C for 25 min, to activate the Rubisco enzyme. To assess the activity, 50 µl of leaf extract was incubated with 1550 μl of EME buffer (100 mM EPPS–NaOH (pH 8.2), 20 mM MgCl_2_, 1 mM Na_2_-EDTA), 40 µl of 10 mM NADH (final concentration 200 µM), 80 µl of 250 mM NaH_12_CO_3_ (final concentration 10 mM), 200 µl of ATP/phosphocreatine (final concentrations 1 mM and 5 mM, respectively), 40 µl of coupling enzymes (carbonic anhydrase 500 U, glyceraldehyde-3-phosphate dehydrogenase 50 U, 3-phosphoglycerate kinase 50 U, triose-phosphate isomerase/glycerol-3-phosphate dehydrogenase 400/40 U), and 40 µl of 25 mM RuBP (final concentration 500 µM), and the decrease of absorbance was monitored at 340 nm for 180 s using a UV–Vis spectrophotometer. The Rubisco carboxylation rate was calculated using the equation:

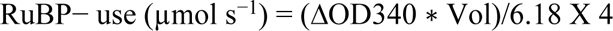

assuming an extinction coefficient of 6.18 µmol ml^−1^ cm^−1^ for NADH oxidation at 340 nm. ΔOD340 is the rate of change of absorbance at 340 nm per second, and Vol is assay volume (1 ml). The equation accounts for the four NADH molecules oxidized per RuBP carboxylated by Rubisco. The activity of Rubisco (*Vcmax*) is calculated as:

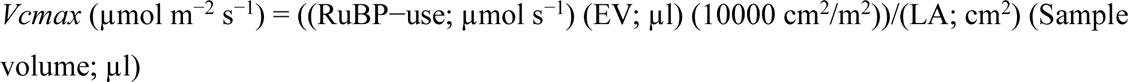

where EV is extraction volume and LA is leaf area.

### Rice protoplast isolation and PEG mediated transformation

Protoplast from rice seedlings were isolated according to Zhang et al., 2011. Seeds of WT, *mpk3 ko* and *actpk1 ko* were surface sterilized and germinated on half-strength MS solid medium. Seven- to ten-day-old seedlings were chopped into 0.5 mm strips in 0.5 M mannitol, incubated for 10 min, and transferred to enzyme solution (0.4 M mannitol, 1.5% Cellulase RS (Yakult), 0.75% Macerozyme (Yakult), 20 mM MES (pH 5.7), 10 mM CaCl_2_, 20 mM KCl, 0.1% BSA) for 3-4 h at 80 rpm at room temperature. The reaction was stopped with equal volume of W5 buffer (154 mM NaCl, 125 mM CaCl_2_, 5 mM KCl, 2 mM MES (pH 5.7)), shaken for 10 s to release protoplasts, and filtered through a 100 µm nylon mesh. Additional 5 ml W5 buffer was added to the strips, strained again and centrifuged at 1500 rpm for 3 min at room temperature. The pellet was washed with 5 mL W5 buffer, resuspended in MMG buffer (4 mM MES (pH 5.7), 0.4 M Mannitol, 15 mM MgCl_2_), and protoplast quality was assessed under a light microscope.

PEG-mediated transient transformation of protoplasts was performed according to Shen et al., (2014). Approximately 100 µL of protoplasts were mixed with 5–10 µg plasmid of RbcS-pGWB5, RbcS^T12D^-pGWB5 and RbcS^T12A^-pGWB5 in a microcentrifuge tube by gentle tapping. Subsequently, 110 µL of PEG solution (40% PEG_4000_ (Fluka), 0.2 M Mannitol, 100 mM CaCl_2_) was added, mixed by tapping, and incubated in the dark at room temperature for 20 min. The solution was diluted with 440 µL W5 buffer, centrifuged at 1500 rpm for 3 min, and the pellet was resuspended in 1 mL WI solution (4 mM MES, 0.5 M mannitol, 20 mM KCl). After 16 h incubation in the dark, transformed protoplasts were analysed using a Leica TCS SP8 AOBS Laser Scanning Microscope (Leica Microsystems) using GFP and chloroplast autofluorescence filters.

For phosphatase inhibitor treatment, protoplasts were exposed to 50nm okadaic acid after 16 h of transfection for 30 mins. Afterwards protoplast were analysed using a Leica TCS SP8 AOBS Laser Scanning Microscope (Leica Microsystems) using GFP and chloroplast autofluorescence filters.

### Pigment content measurement

Pigment content were measured according to procedure described by Porra et al., (1989). Leaf samples (1 g) were harvested post photosynthetic parameters measurements, and chlorophyll and total carotenoid was extracted in 80% ice-cold acetone (aqueous). Chlorophyll and carotenoid contents were measured spectrophotometrically and calculated according to Arnon, 1949.

### Statistical analysis

Student t-test and one-way analysis of variance (ANOVA) were performed to determine significant differences indicated as *P<0.05, **P< 0.005, and ***P< 0.0005. Details of statistical analysis and sample sizes in each experiment are mentioned in the figure legends. ImageJ was used to quantify the intensities of bands in the immunoblots.

## Locus Ids of gene

MPK3 (LOC_Os03g17700); MPK6 (LOC_Os06g06090); ACTPK1 (LOC_Os02g02780); RbcS3 (LOC_Os12g17600)

## Supplemental figure and table legends

**Fig S1.** Photosynthetic pigments and parameters in WT, *MPK3 OE* (*MPK3-23*, *MPK3-24*) and *mpk3 ko* (*mpk3 ko1*, *mpk3 ko2*) lines.

**Fig. S2** Localisation of MPK3 and interaction and phosphorylation of RbcS with MPK3 and MPK6.

**Fig. S3** ACTPK1 is a Raf-like MAPKKK exhibiting homology and sequence similarity with *At*STY8, *At*STY17 and *At*STY46, and contains multiple MAPK phosphorylation motifs.

**Fig. S4** Lysine at 332 and 428 are crucial for trans- and auto-phosphorylation activity of ACTPK1 and ACTPK1 do not phosphorylate MPK3.

**Fig. S5** ACTPK1 phosphorylates RbcS at Thr12.

**Fig. S6** Localization of RbcS-GFP in *mpk3 ko* and *actpk1 ko* protoplasts.

**Fig. S7** Development of *ACTPK1 OE* and *actpk1 ko* lines.

**Fig. S8** Intercellular CO_2_ concentration in flag leaf of WT, *ACTPK1 OE* and *actpk1 ko* plants.

**Fig. S9** ACTPK1 regulates vegetative growth, flag leaf angle, flowering time and seed architecture.

**Supplemental Table 1**: Table containing the list of primers used in this study.

## Acknowledgments

SJ and RB are thankful to the Department of Biotechnology (DBT) for the fellowship. GB, MB and PS are thankful to the Council of Scientific and Industrial Research (CSIR) for the fellowship. AKS thanks Sir JC Bose Fellowship from the Anusandhan National Research Foundation, Department of Science and Technology, Government of India (File no.: JCB/2020/000041). The work is supported by the grant from the Anusandhan National Research Foundation, Department of Science and Technology, Government of India (File No.: CRG/2023/001751). The authors also acknowledge the Gene Functional Analysis Platform for Crops, Confocal Facility, DNA sequencing facility, Radioisotope facility and the Central Instrumentation Facility of NIPGR, New Delhi, India. The authors declare no conflict of interest.

## Authors’ contribution

SJ and AKS planned the study and designed the experiments. SJ and RB generated and screened transgenic plants and recorded the photosynthetic parameters. SJ and GB carried out the molecular biology experiments. SJ, MB and MM recorded and analysed the phenotypic traits of transgenic plants. PS screened the *mpk3actpk1* double knockout plants. SJ and AKS analysed data and wrote the manuscript. SJ wrote the initial draft and AKS finalise the draft. All authors read and approved the final manuscript.

