## Supplementary material for "Cytosolic MAPK signaling gates chloroplast protein import and photosynthetic capacity": Supplimental File

Supplemental File

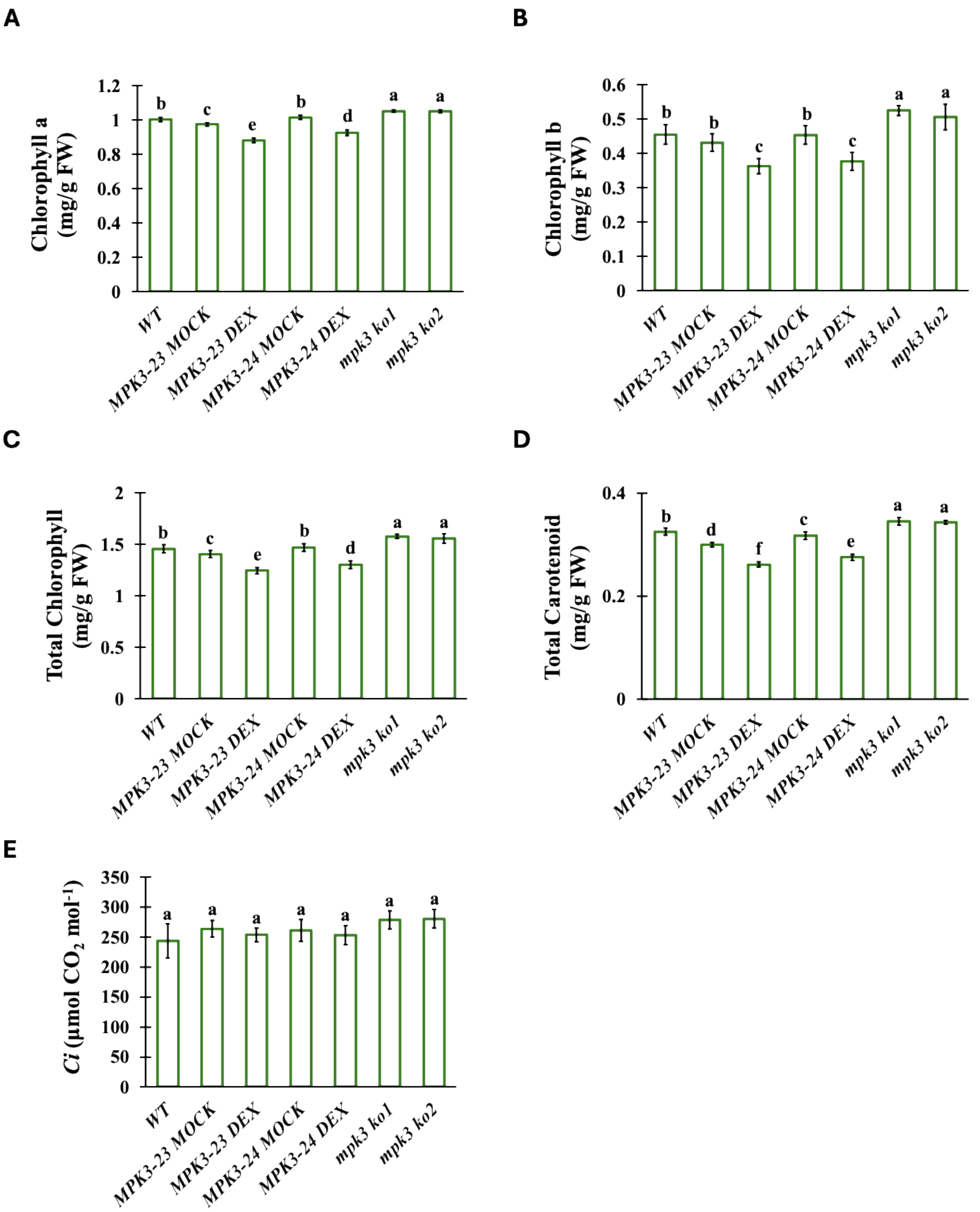


**Fig S1. Photosynthetic pigments and parameters in WT, *MPK3 OE* (*MPK3-23*, *MPK3-24*) and *mpk3 ko* (*mpk3 ko1*, *mpk3 ko2*) lines. (A-D)** Bar graph showing the Chlorophyll a **(A)**, Chlorophyll b **(B)**, total chlorophyll **(C)**, and total carotenoid content **(D)** in WT, *MPK3 OE* (*MPK3-23*, *MPK3-24*) and *mpk3 ko* (*mpk3 ko1*, *mpk3 ko2*) lines. Each value represents mean ± SD, where n = 15 data points from different plants. Different letters indicate statistical significance according to one-way ANOVA followed by post-hoc Tukey HSD calculation at $P < 0.05$. **(E)** Bar graph representing intercellular $CO_2$ concentration at ambient $CO_2$ (400 µmol $mol^{-1}$) in WT, *MPK3 OE* (*MPK3-23*, *MPK3-24*) and *mpk3 ko* (*mpk3 ko1*, *mpk3 ko2) lines. Each value represents mean ± SD, where n = 5 data points from different plants. Different letters indicate statistical significance according to one-way ANOVA followed by post-hoc Tukey HSD calculation at $P < 0.05$. MOCK, dimethyl sulfoxide treated (DMSO) plants; DEX, dexamethasone treated plants. 1 µM DMSO and DEX sprayed on *MPK3 OE* leaves, 24 h prior to taking measurements. All the photosynthetic measurements were taken from flag leaf at heading stage using LiCOR6800, between 8:00 am to 11:00 am.

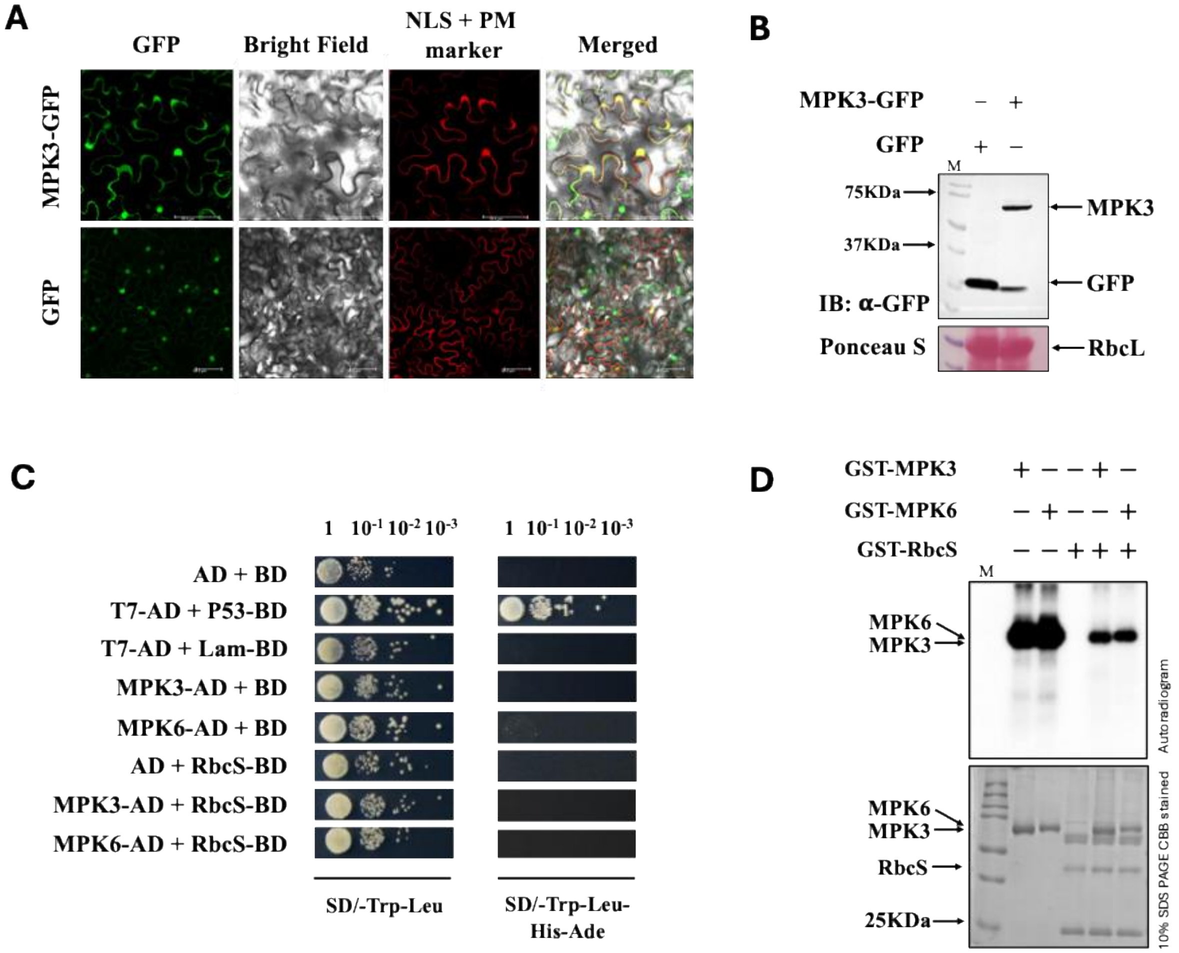


**Fig. S2 Localisation of MPK3 and interaction and phosphorylation of RbcS with MPK3 and MPK6. (A)** Confocal images depicting the localisation of MPK3 to nucleus and cytosol. Empty-GFP localisation served as negative control. Scale Bar, 25 $\mu$m. **(B)** Western blots showing presence of MPK3 and GFP in confocal localisation samples of (A). Ponceau stained blot shows RbcL protein of *N. benthamiana*. **(C)** MPK3 and MPK6 does not interact with RbcS in yeast cells. Yeast cells were cultured on SD/-Trp-Leu and SD/-Trp-Leu-His-Ade media. AD, GAL4 activation domain; BD, GAL4 DNA-binding domain; SD, synthetic defined. **(D)** *In vitro* kinase assay showing GST-MPK3 and GST-MPK6 does not phosphorylate GST-RbcS. CBB stained 10% SDS PAA gel shows loaded proteins.

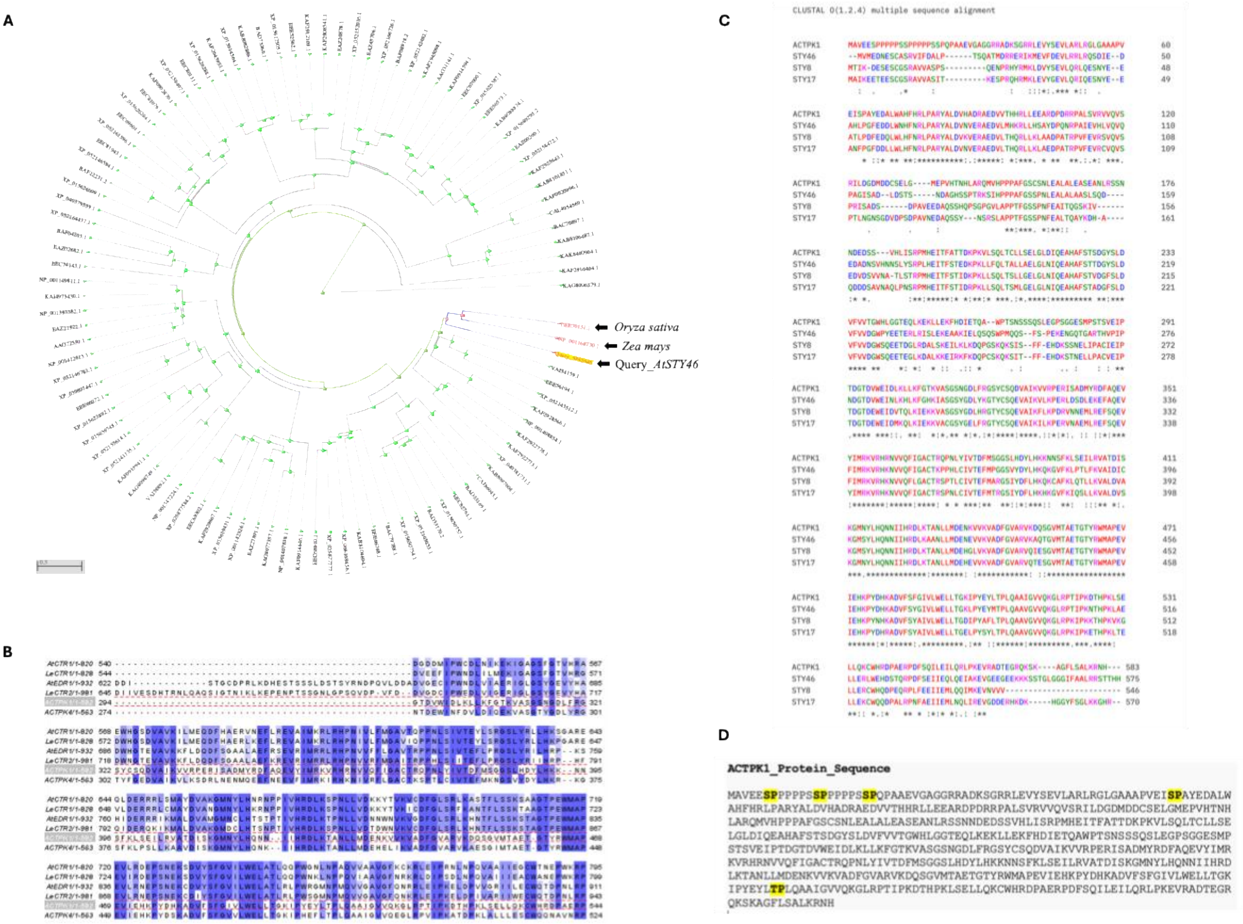


**Fig. S3 ACTPK1 is a Raf-like MAPKKK in rice exhibiting homology and sequence similarity with *At*STY8, *At*STY17 and *At*STY46, and contains multiple MAPK phosphorylation motifs.** (A) Distant tree showing the homologues of *At*STY46. Accession numbers written in red are the closest homologue of *Arabidopsis* STY46 in *Zea mays* and *Oryza sativa*. Query sequence is taken as the AtSTY46 protein sequence against nr_cluster_seq database (NCBI) and the tree was produced by BLAST pairwise alignment. **(B)** Sequence Alignment of ACTPK1 Kinase domain with identified Plant Raf like MAPKKKs Kinase domain. Shaded boxes indicate identical residues. Prefixes on protein names indicate species of origin. *At*, *Arabidopsis thaliana*; *Le, Solanum lycopersicum var. lycopersicum.* Alignment was constructed using Clustal Ω program and analysed using Jalview software. **(C)** Sequence Alignment of ACTPK1 with AtSTY8, AtSTY17 and AtSTY46. Alignment was constructed using Clustal 0(1.2.4). **(D)** Protein sequence of ACTPK1 showing multiple MAPKs phosphorylation motifs (S/T followed by P). MAPKs phosphorylation motifs were highlighted in yellow colour.

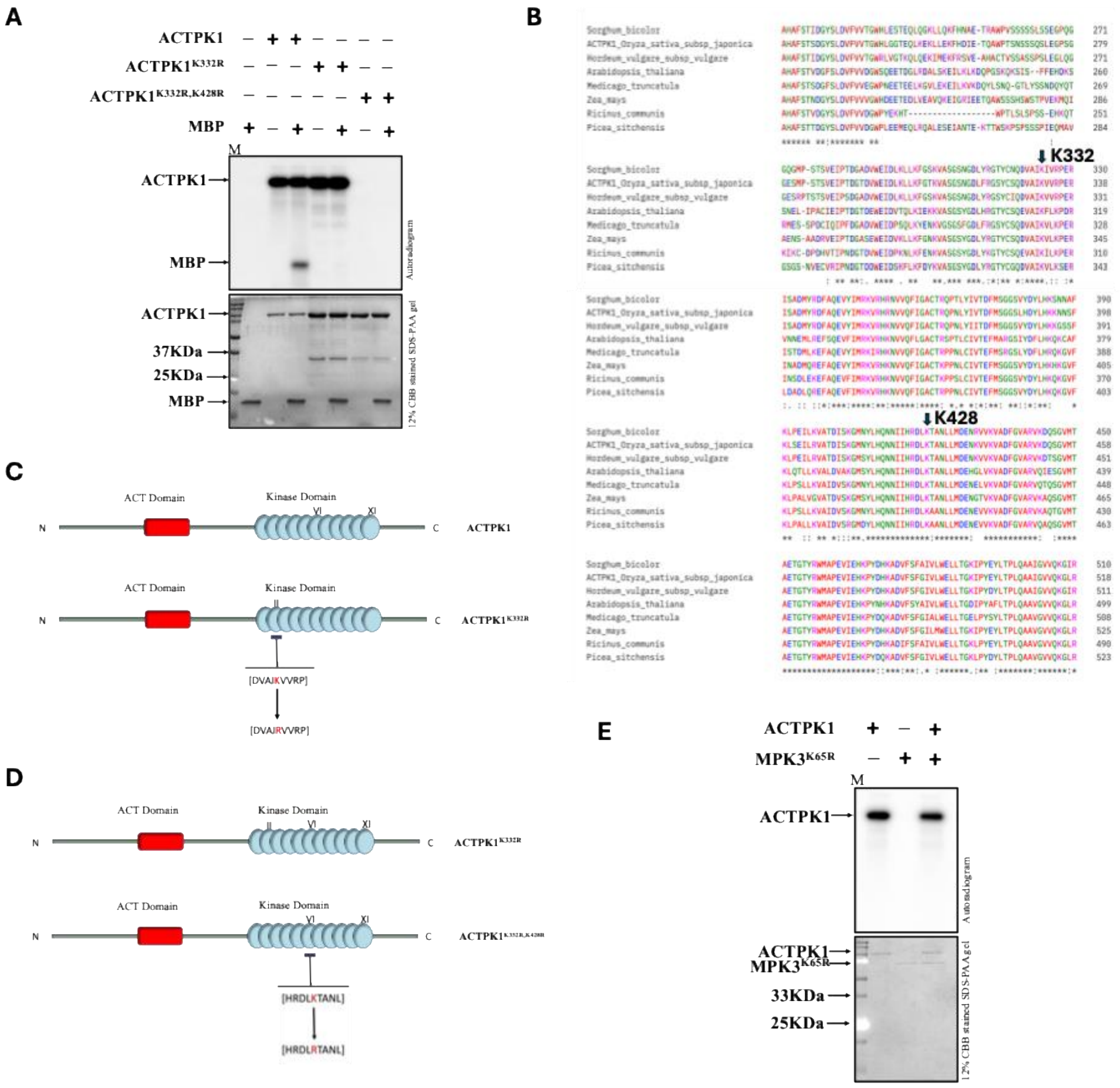


**Fig. S4 Lysine at 332 and 428 are crucial for trans- and auto-phosphorylation activity of ACTPK1 and ACTPK1 do not phosphorylate MPK3. (A)** *In vitro* phosphorylation assay depicting ACTPK1 showed auto- and trans-phosphorylation, ACTPK1$^{K332R}$ showed only autophosphorylation and ACTPK1$^{K332R,K428R}$ does not showed auto- or trans-phosphorylation. MBP was used as a substrate for ACTPK1, ACTPK1-GST, ACTPK1$^{K332R}$-GST, ACTPK1$^{K332R,K428R}$-GST**.** CBB stained 12% SDS PAA gel shows loaded proteins. **(B)** Diagrammatic representation showing the mutation created for generating ACTPK1$^{K332R}$. **(C)** Diagrammatic representation showing the mutation created for generating ACTPK1$^{K332R,K428R}$. **(D)** Sequence alignment of ACTPK1 kinase domain with its homologues from different plants showing the conservation of lysine at 332 (showed by black arrow followed by K332) and lysine at 428 (showed by black arrow followed by K428). Alignment was constructed using Clustal 0(1.2.4) **(E)** *In vitro* phosphorylation assay depicting ACTPK1 do not phosphorylates kinase dead MPK3$^{K65R}$-GST. CBB stained 12% SDS PAA gel shows loaded proteins.

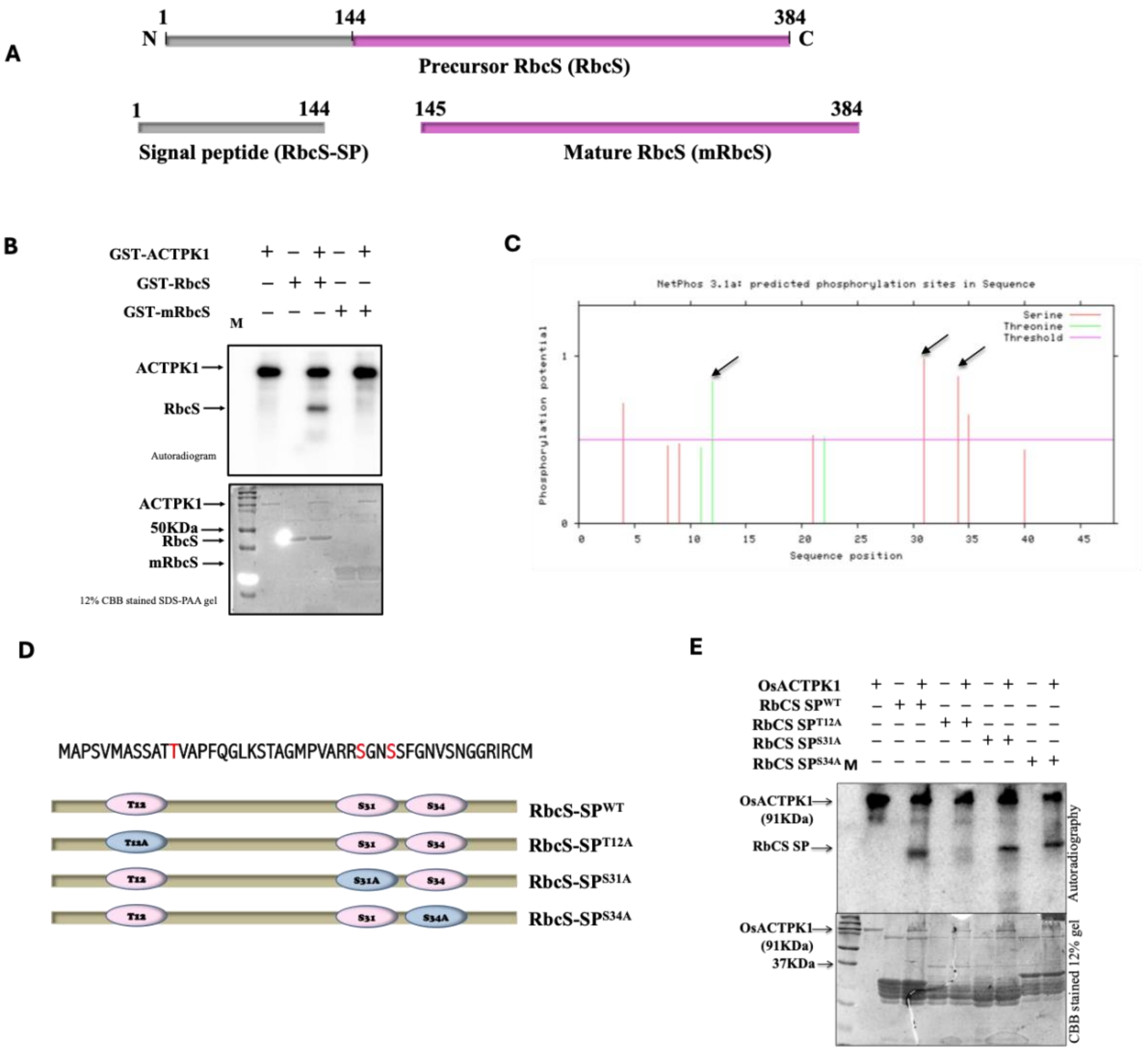


**Fig. S5 ACTPK1 phosphorylates RbcS at Thr12. (A)** Diagrammatic picture representing the truncation of precursor RbcS protein into signal peptide (RbcS-SP) and mature RbcS (mRbcS). **(B)** *In vitro* kinase assay showing ACTPK1 phosphorylates precursor RbcS (GST-RbcS) but not mature RbcS (mRbcS). CBB stained 12% SDS PAA gel shows loaded proteins. **(C)** NetPhos3.1 results showing the phosphorylation potential of different amino acids of RbcS. Black arrow indicates the amino acids having high phosphorylation potential score. **(D)** Pictorial representation of all the RbcS mutant constructs designed through site-directed mutagenesis. **(E)** *In vitro* kinase assay depicting Thr at 12 is the site of ACTPK1 phosphorylation. CBB stained 12% SDS PAA gel shows loaded proteins.

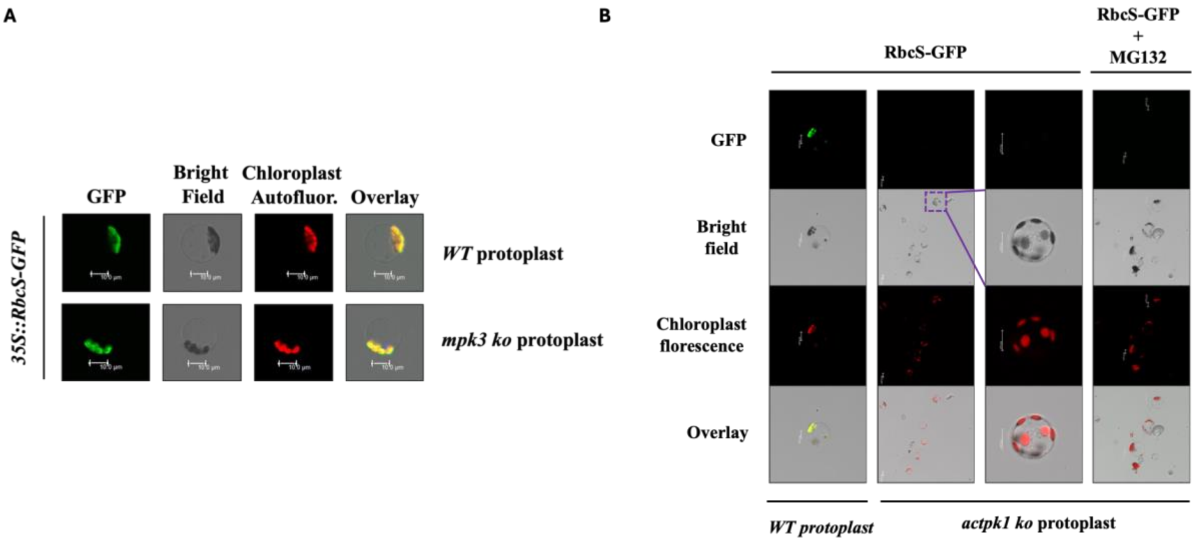


**Fig. S6 Localization of RbcS-GFP in *mpk3 ko* and *actpk1 ko* protoplasts. (A)** Localization of RbcS in protoplast isolated from WT and *mpk3 ko* rice seedlings. Scale Bar, 10 µm. **(B)** Localization of RbcS in protoplast isolated from WT and *actpk1 ko* rice seedlings. Scale Bar, 10 µm. Third panel is the enlarged image of protoplast present in purple box of second panel. MG132 treatment was given for 4h.

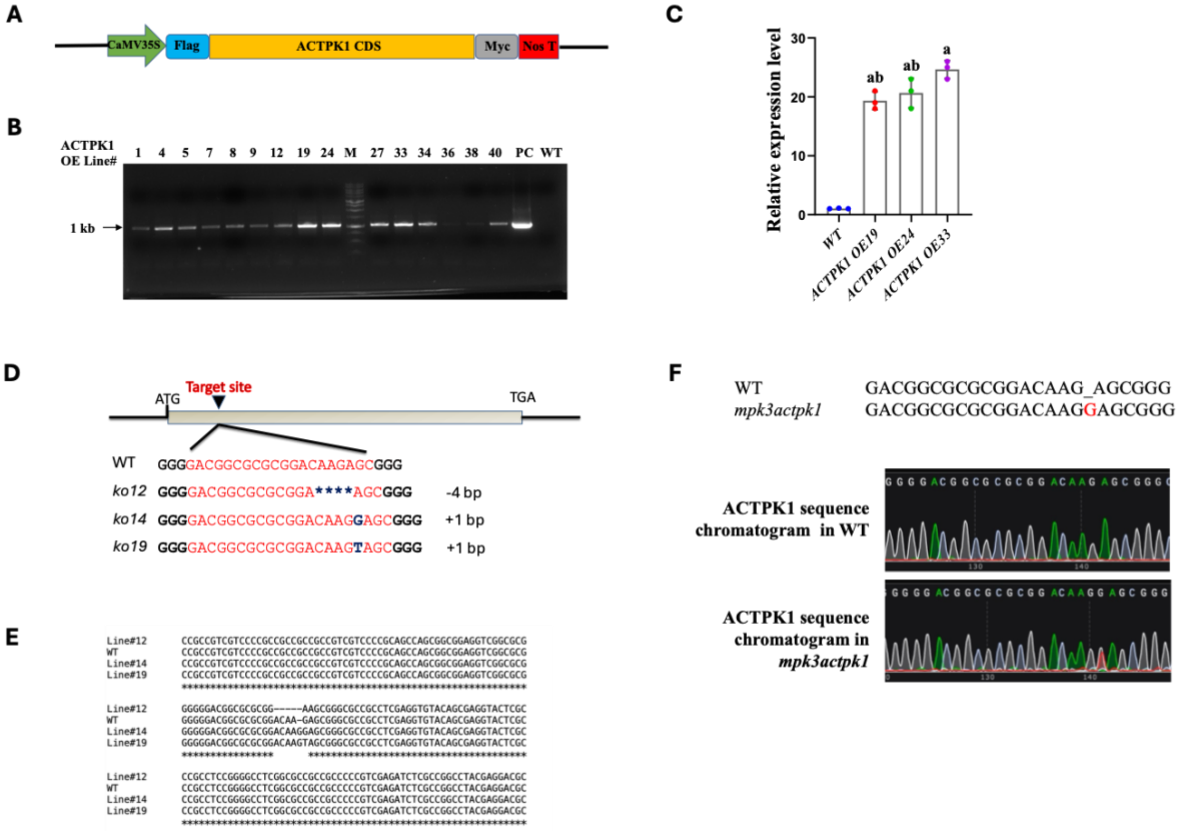


**Fig. S7 Development of *ACTPK1 OE* and *actpk1 ko* lines. (A)** Diagrammatic representation of the construct used for generating *ACTPK1 OE* plants **(B)** Nucleic acid gel image showing Semi-quantitative Real-time PCR analysis of *ACTPK1* expression in overexpression lines compared with WT plant. **(C)** Bar graph showing relative expression level of ACTPK1 in *ACTPK1 OE19, ACTPK1 OE24* and *ACTPK1 OE33* lines. Each value represents mean ± SD, where n = 3 biological replicates. Different letters indicate statistical significance according to one-way ANOVA followed by post-hoc Tukey HSD calculation at $P < 0.05$. **(D)** Diagrammatic representation of target site and sgRNA sequence and mutations observed in *actpk1 ko 12, actpk1 ko 14, and actpk1 ko 19* lines **(E)** Sequence alignment of different mutation identified from *actpk1 ko 12, actpk1 ko 14, and actpk1 ko 19* lines using clustal omega software. **(F)** Chromatogram of WT and *mpk3actpk1* showing insertion of G in coding region of ACTPK1 in *mpk3actpk1 ko* line. Inserted base is depicting in red colour. Knockout mutation in ACTPK1 was generated in the background of *mpk3 ko2.*

**A**

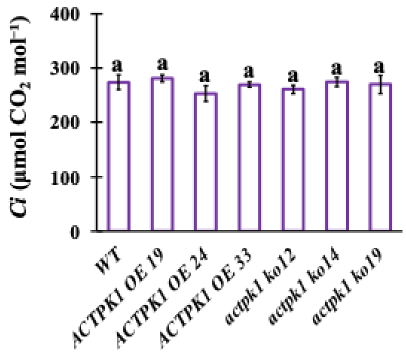


**Fig. S8 Intercellular $CO_2$ concentration in flag leaf of WT, *ACTPK1 OE* and *actpk1 ko* plants. (A)** Bar graph showing intercellular $CO_2$ concentration at ambient $CO_2$ (400 µmol mol$^{-1}$) in WT, *ACTPK1 OE* (ACTPK1 OE19, ACTPK1 OE24 and ACTPK1 OE33) and *actpk1 ko* (*actpk1 ko 12, actpk1 ko 14, and actpk1 ko 19*) lines. Each value represents mean ± SD, where n = 5 data points from different plants. Different letters indicate statistical significance according to one-way ANOVA followed by post-hoc Tukey HSD calculation at $P < 0.05$. All the photosynthetic measurements were taken from flag leaf at heading stage using LiCOR6800, between 8:00 am to 11:00 am.

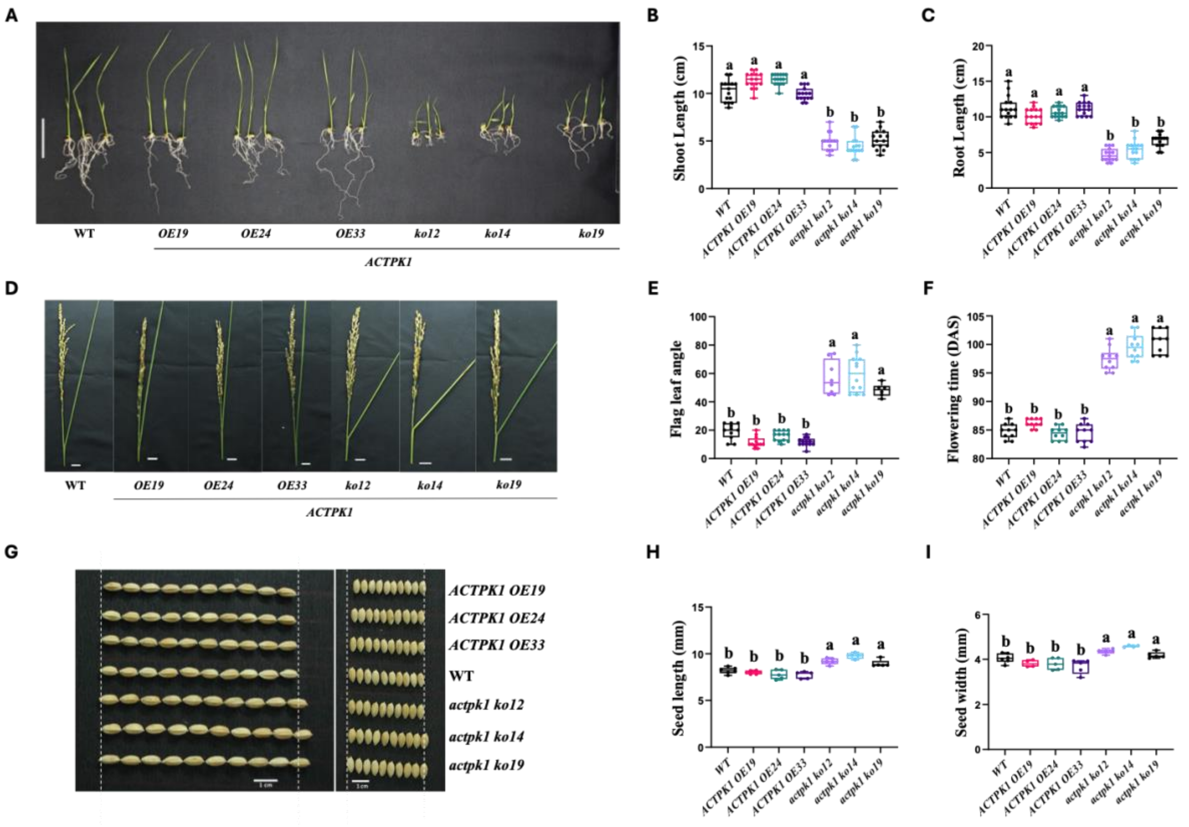


**Fig. S9 ACTPK1 regulates vegetative growth, flag leaf angle, flowering time and seed architecture. (A)** Representative images of WT, *ACTPK1 OE* and *actpk1 ko* seedlings. Scale Bar, 10 cm. **(B-C)** Bar graph showing shoot length (B) and root length (C) of 10 day old seedling of WT, *ACTPK1 OE* (ACTPK1 OE19, ACTPK1 OE24 and ACTPK1 OE33) and *actpk1 ko* (*actpk1 ko 12, actpk1 ko 14, and actpk1 ko 19*). Each value represents mean ± SD, where n = 10 data points. Different letters indicate statistical significance according to one-way ANOVA followed by post-hoc Tukey HSD calculation at $P < 0.05$. **(D)** Representative images depicting the flag leaf angle of WT, *ACTPK1 OE* and *actpk1 ko* plants**.** Scale Bar, 1 cm. **(E)** Bar graph depicting flag leaf angle of WT, *ACTPK1 OE* and *actpk1 ko* plants. Each value represents mean ± SD, where n = 10 data points. Different letters indicate statistical significance according to one-way ANOVA followed by post-hoc Tukey HSD calculation at $P < 0.05$. **(F)** Bar graph showing flowering time of WT, *ACTPK1 OE* and *actpk1 ko* plants. Each value represents mean ± SD, where n = 10 data points. Different letters indicate statistical significance according to one-way ANOVA followed by post-hoc Tukey HSD calculation at $P < 0.05$. **(G)** Representative pictures depicting the seed length and seed width of WT, (ACTPK1 OE19, ACTPK1 OE24 and ACTPK1 OE33) and *actpk1 ko* (*actpk1 ko 12, actpk1 ko 14, and actpk1 ko 19*) plants. Scale Bar, 1 cm. **(H-I)** Bar graph depicting the seed length (H) and seed width (I) of WT, *ACTPK1 OE* and *actpk1 ko* lines. Each value represents mean ± SD, where n = ~1000 data points from at least 5 different plants. Different letters indicate statistical significance according to one-way ANOVA followed by post-hoc Tukey HSD calculation at $P < 0.05$.

**Supplemental Table1**: Table containing the list of primers used in this study.

| **Primer Name** | **Sequence (5`-3`)** |
|---|---|
| **Primers used for Gateway entry cloning** | |
| MPK3 pENTR F | CACCATGGACGGGGCGCCGGTGGCGG |
| MPK3 pENTR R | GTACCGGAAGTTTGGGTTCATCTC |
| MPK6 pENTR F | CACCATGGACGCCGGGGCGCAGCCGTC |
| MPK6 pENTR R | CTGGTAATCAGGGTTGAACGCAAG |
| ACTPK1 pENTR F | CACCATGGCCGTGGAGGAGTCGCC |
| ACTPK1 pENTR R | GTGGTTTCTCTTCAATGCTG |
| RbcS pENTR F | CACCATGGCCCCCTCCGTGATG |
| RbcS pENTR R | GTTGCCACCAGACTCCTCGC |
| **Primers used for Protein Expression** | |
| MPK3 pGEX F | GGATCCATGGACGGGGCGCCGGTGGC |
| MPK3 pGEX R | CTCGAGCGGGTACCGGAAGTTTGGGT |
| MPK6 pGEX F | GGATCCATGGACGCCGGGGCGCAGCCG |
| MPK6 pGEX R | CTCGAGCTGGTAATCAGGGTTGAAC |
| ACTPK1 pGEX F | CACCGAATTCCCATGGCCGTGGAGGAGTCGC |
| ACTPK1 pGEX R | CCGTCGACGTGGTTTCTCTTCAATGC |
| RbcS pGEX F | TTGGATCCATGGCCCCCTCCGTGATG |
| RbcS pGEX R | TTGTCGACGCCACCAGACTCCTCGCA |
| RbcS-SP pGEX F | TTGGATCCATGGCCCCCTCCGTGATG |
| RbcS-SP pGEX R | TTGTCGACCCTGATCCTGCCGCCATTGC |
| mRbcS pGEX F | CACCCAGGTGTGGCCGATTGAGGGCAT |
| mRbcS pGEX R | TTGTCGACGCCACCAGACTCCTCGCA |
| **Primers used for Yeast-2 Hybrid assay** | |
| ACTPK1 AD/BD F | AAGAATTCATGGCCGTGGAGGAGT |
| ACTPK1 AD/BD R | CCGGATCCGTGGTTTCTCTTCAATGC |

| | |
|---|---|
| MPK3 AD/BD F | CATATGATGGACGGGGCGCCGGT |
| MPK3 AD/BD R | GGATCCGTACCGGAAGTTTGGGT |
| MPK6 AD/BD F | GGATCCGGATGGACGCCGGGGCGCAGC |
| MPK6 AD/BD F | GGATCCGCTACTGGTAATCAGGGTTG |
| **Primers used for developing and screening transgenic plants** | |
| Cas9 F | AACAGCCGCGAGAGAATGAA |
| Cas9 R | TCGGCCTTGGTCAGATTGTC |
| ACTPK1 sgRNA F | GGCAGACGGCGCGCGGACAAGAGC |
| ACTPK1 sgRNA R | AAACGCTCTTGTCCGCGCGCCGTC |
| ACTPK1 Crispr_Seq F | TCTCCTCTCTCGTCTCGTG |
| ACTPK1 Crispr_Seq R | CATGGATGGATCTCTCTCA |
| CaMV 35S_F | CCACCATGTTGGCATGGAGTCAAAGATTCAAATAG AG |
| CaMV 35S_R | GAATTCAGTCCCCCGTGTTCTCTCCAAATG |
| FLAG_F | GGATCCATGGACTATAAGGACCACGAC |
| **SDM** | |
| ACTPK1 K332R F | AGTCAGGATGTAGCTATTAGAGTAGTGAGACCTGAACGC |
| ACTPK1 K332R R | GCGTTCAGGTCTCACTACTCTAATAGCTACATCCTGACT |
| ACTPK1 K428R F | CATAATTCACCGTGATCTCAGGACAGCGAATCTTCTTATGG |
| ACTPK1 K428R F | CCATAAGAAGATTCGCTGTCCTGAGATCACGGTGAATTATG |
| MPK3 K65R F | GATGGTGGCGATACGCAAGATCGCCAACGC |
| MPK3 K65R R | GCGTTGGCGATCTTGCGTATCGCCACCATC |
| RbcS T12A F | CACCATGGCCCCCTCCGTGATGGCGTCGTCGGCCACCGCCG TCGCTCCCTTC |
| RbcS T12A R | GTTGCCACCAGACTCCTCGC |
| RbcS S31A F | CACCATGGCCCCCTCCGTGATG |
| RbcS S31A R | CATGCACCTGATCCTGCCGCCATTGCTGACGTTGCCGAAGC TGGCGTTGCCGGAGCGGC |
| RbcS S34A F | CACCATGGCCCCCTCCGTGATG |
| RbcS S34A R | CATGCACCTGATCCTGCCGCCATTGCTGACGTTGCCGAAGC TGGAGTTGCCGGCGCGGC |

| | |
|---|---|
| RbcS T12D F | CACCATGGCCCCCTCCGTGATGGCGTCGTCGGCCACCGACGTCGCTCCCTTC |
| RbcS T12D R | GTTGCCACCAGACTCCTCGC |
| **Primers used for RT-PCR** | |
| ACTPK1 RT F | CCGTGATCTCAAGACAGCGA |
| ACTPK1 RT R | CTCAGGTGCCATCCAACGAT |
| RbcS RT F | AACCACAGATCCCCCGGATA |
| RbcS RT R | CCTGCCTGACGTTGTCGAA |
| Actin7 RT F | CGGTGTGATGGTTGGTATGG |
| Actin7 RT R | GCCTCAGTCAGCAACACAGG |
| UBQ5 RT F | ACCACTTCGACCGCCACTACT |
| UBQ5 RT R | ACGCCTAAGCCTGCTGGTT |
